# Inferring state-dependent functional circuit motifs using higher-order interactions analysis

**DOI:** 10.64898/2026.03.09.710501

**Authors:** Safura Rashid Shomali, Seyyed Nader Rasuli, Hideaki Shimazaki, Sadra Sadeh

## Abstract

Analysing higher-order interactions among simultaneously recorded neurons can provide crucial insights into neural network dynamics. Recent technological advances have enabled large-scale, long-term neuronal recordings, but analysis of such datasets often relies on simpler statistics due to computational and statistical challenges in assessing higher-order interactions. Here, we developed CHOIR, an efficient and reliable method for calculating higher-order interactions from large-scale neuronal recordings. We then used the inferred HOIs to uncover the underlying functional connectivity, differentiating between connectivity motifs in the space of pairwise and triplet-wise interactions. We found that this approach could successfully distinguish stationary and running states, sleep and awake states, and neuronal ensembles with distinct activity patterns in mice. Furthermore, we identified potential circuit architectures underlying different higher-order interactions, which we confirmed through simulations of large-scale spiking networks with specific subnetwork connectivity. Applying CHOIR to a causal manipulation dataset further confirmed the role of lateral inhibition, a key inhibitory motif, in generating specific HOI patterns. Our work provides a systematic analysis of higher-order interactions in diverse datasets and suggests that HOIs can reveal circuit motifs underlying neural dynamics across brain areas and brain states.

## Introduction

Neuronal networks display non-random connectivity patterns, with certain motifs occurring more frequently than expected by chance ^1–3^. Deciphering these connectivity motifs from neural activities is challenging, but essential to understanding brain circuit dynamics and function. Although connectomics provides detailed maps of anatomical connections ^4,5^, structural connectivity alone cannot predict which neurons will be active together during behaviour, as the same anatomical circuit can give rise to different functional states ^6,7^. Functional connectivity therefore reveals how the brain dynamically engages specific pathways across different behavioural states, capturing information that static anatomical connectivity alone cannot provide ^6,7^.

Recent technological advances have enabled simultaneous recording of large neuronal populations over extended periods of time, providing a unique opportunity to analyse neural dynamics ^8–10^. However, we still cannot record from all neurons and their inputs simultaneously to reveal the effective connectivity (but see ^6,7^), which makes the analysis of neuronal interactions crucial for inferring functional microcircuits from observed activity patterns.

Network interactions generate emergent dynamics that shape collective behaviour ^11–13^. Several approaches using information theory ^14^, information geometric measure of interactions ^15–18^or computational topology ^19–21^have demonstrated that the brain exhibits higher-order interactions (HOIs, i.e., interactions among three neurons or more) that crucially shape the complexity of neural dynamics ^22–26^. Pairwise and higher-order interactions (HOIs) between neurons have been observed ^22,27–30^and contribute significantly to stimulus encoding and population coding ^19,23,31–37^. HOIs have been associated with cognitive function including expectation ^36^, perceptual accuracy enhancement ^38^, decision-making ^39^, and memory-related behaviour ^40^. Interestingly, recent experimental work have shown that using HOIs, neuronal activities can be differentiated in amnestic and wild-type animals ^40^, healthy and clinical groups ^41–43^, patients with different levels of consciousness ^44^ and aging ^45^. These studies raise the possibility that HOIs can be used to distinguish between different functional states, and between functional and dysfunctional neural circuits.

HOIs can reflect coordinated activity arising from shared circuit motifs and common inputs among neurons ^46,47^. Since cell assemblies are precisely defined by such coordinated activity patterns, ignoring HOIs may mask the true functional architecture of cell assemblies ^30,48,49^. Recent theoretical work has demonstrated that the sign and magnitude of HOIs are jointly determined by the type of shared input and the underlying network architecture, with distinct excitatory and inhibitory motifs occupying non-overlapping regions in the triple-wise versus pairwise interaction plane, enabling unambiguous identification of hidden circuit motifs directly from observed population statistics ^50^. However, the relationship between HOIs and various circuit motifs requires further investigation.

Two major challenges have limited HOI analysis: reliably estimating interactions from neural activity recordings, and the computational cost of testing all possible neuron combinations ^49,51^. To address these, here we have developed CHOIR (Circuit motifs from Higher-Order Interactions in neural Recordings), a computationally efficient approach for measuring HOIs in large-scale neural recordings. In the following, we first showed that our method can reliably and efficiently extract statistically significant interactions from large-scale spiking activity datasets, overcoming the computational challenges associated with calculating and interpreting HOIs. We tested this on multiple datasets in mice, including Neuropixels recordings from Allen Brain Observatory datasets ^52^, extracellular recording of sleep and awake data ^53^, and two-photon calcium imaging during visual stimulation ^54^. Next, we showed that analysing the inferred interactions in the space of pairwise and triple-wise interactions ^50^ can reveal insights into the underlying functional circuit motifs, which cannot be inferred from pairwise interactions alone. We then investigated whether and how HOIs could distinguish different behavioural states of the animals. To this end, we analysed distinct patterns of HOIs to uncover different circuit motifs during stationary versus running, and in sleep versus awake, states.

## Results

### Inferring higher-order interactions from spiking activity

We quantified neuronal interactions by modelling individual and joint spiking activities using exponential family distributions, which capture both independent firing patterns and coordinated activity between neurons (Methods) ^15,27,55^. Exponential family distributions arise naturally from the maximum entropy principle, which provides a principled way of modelling interactions while making minimal assumptions about the underlying mechanisms ^13,15,27,47,55^. The distribution’s parameters (Eq. 2 and Eq. 4 in Methods) correspond directly to pairwise and triple-wise interactions. This allows the model to capture higher-order interactions while keeping the same mathematical framework across all orders of interactions ^56^.

Using this approach, we calculated neuronal pairwise and triple-wise interactions between simultaneously recorded neurons. For every combination of three neurons (Fig. 1a), we calculated pairwise interactions between each pair of neurons (Eq. 2) and triple-wise interactions between all three neurons (Eq. 4, Fig. 1b). It is difficult to estimate significant HOIs from experimental data, especially from sparse spiking activity because of the limited sample size. To make sure that the estimated HOIs are indeed statistically significant, we developed HOI metrics which are controlled for random permutations (Methods, and Fig. 1). To this end, we compared the calculated HOI estimates for each triplet with the distributions obtained from the shuffled spike trains (Methods; Fig. 1b). Spike trains were shuffled multiple times (i.e., 100 times per spike trains), and HOIs for each triplet were calculated in the same way. Statistically significant interactions were computed by comparing the actual HOIs with the mean and variance of the distribution of HOIs obtained from this analysis (Methods, Fig. 1b, bottom row).

**Fig 1.**
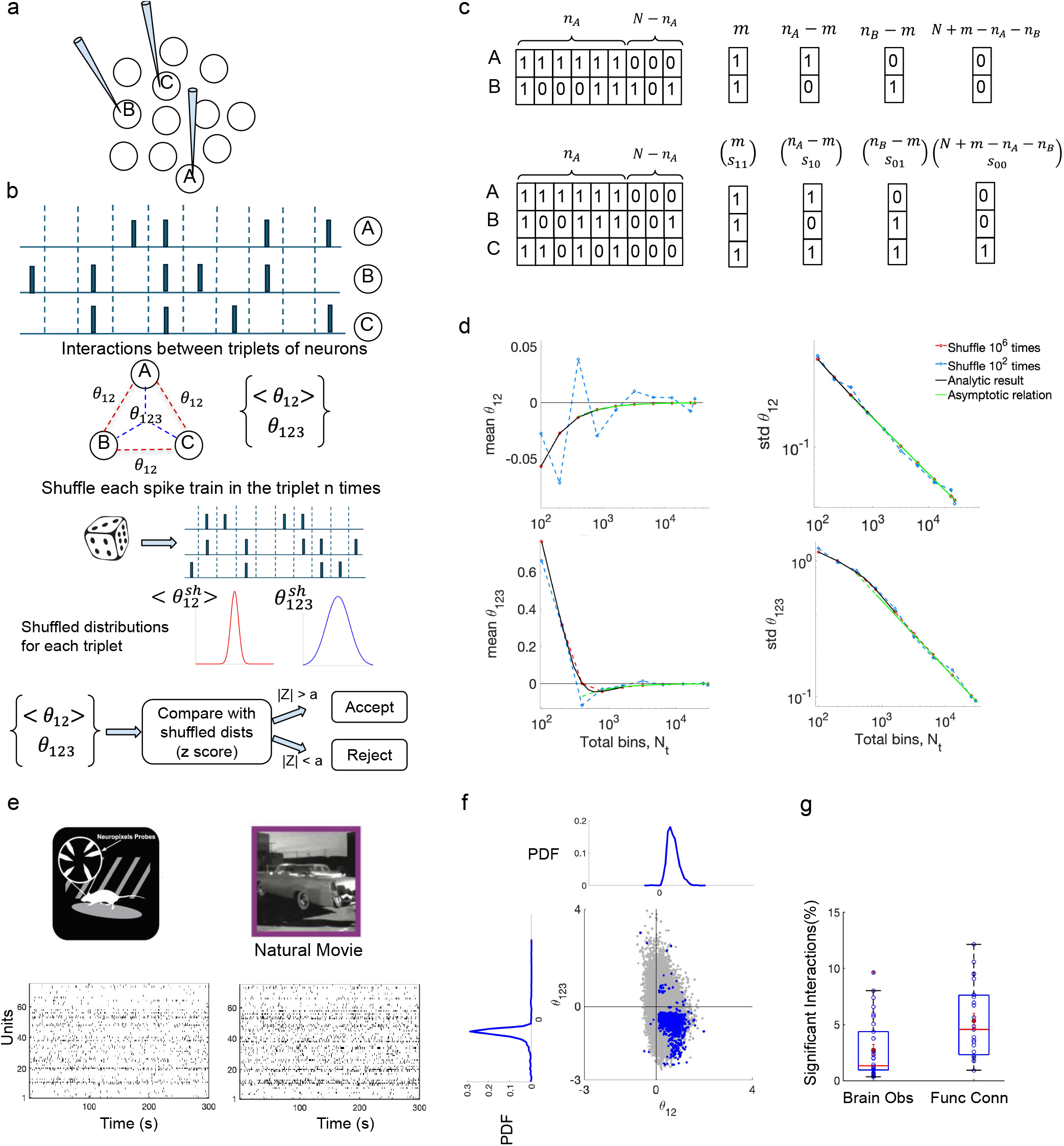
Detecting significant interactions among simultaneously recorded neurons. **a**. Pairwise and triple-wise interactions are calculated among each triplet of neurons (A, B and C) chosen from the simultaneous recordings of neuronal activity. **b** For each triplet, the average pairwise interaction over all possible pairs and the triple-wise interaction among the trio are calculated. The spike trains are then shuffled, and the pairwise and triple-wise interactions are calculated from the shuffled spike trains. Significant interactions are identified by comparing the observed interactions (⟨ θ_12_⟩, θ_123_) to the shuffled distributions and retaining those that exceed significance threshold (Methods). **c** Analytical shuffling of spike trains for two (top) and three (bottom) neurons. In each case, we calculated the number of patterns based on neuron’s firing rates. For two neurons active in *n*_*A*_ and *n*_*B*_ bins respectively (*n*_*B*_ ≤ *n*_*A*_) out of *N*_*t*_ total bins, the joint probabilities are: *P*_11_ = *m*/*N*_*t*_, *P*_10_ = (*n*_*A*_− *m*)/*N*_*t*_, *P*_01_ = (*n*_*B*_ *−m*)/*N*_*t*_, and *P*_00_ = (*N*_*t*_ + *m −n*_*A*_ *−n*_*B*_)/*N*_*t*_, where *m* is the number of co-active bins. From these pattern probabilities, the mean and standard deviation of θ_12_ and θ_123_ distributions across all combinations can be calculated (Methods). **d** Comparison of analytical shuffling with simulation-based shuffled distributions. Mean and standard deviation of null distributions of pairwise and triple-wise interactions are plotted as a function of total bin count (*N*_*t*_) for two neurons (top) and three neurons (bottom). The baseline parameters are *n*_1_ = 25, *n*_2_ = 20, *n*_3_ = 15, *N*_*t*_ = 100, with all spike counts scaled proportionally as *N*_*t*_ increases. The analytic solution (exact as black line and asymptotic relation as green line) is compared with simulations of 100 shuffles (dashed blue) and 10^6^ shuffles (dashed red). **e** We analysed datasets from Allen Brain Observatory, in which awake behaving, head-fixed mice (left) respond to natural movies (example image frame shown on the right). The neural spiking activity of neurons is measured by Neuropixels probes and the behavioural state of the animal is monitored (as assayed by parameters like running speed on the wheel and changes in pupil size). Bottom, example raster plots depicting the spiking activity of units in the primary visual cortex (V1) of a mouse responding to natural movie stimuli. The natural movie (30 second long) is presented 10 times across two blocks which are separated in time (Methods). The data is from Brain Observatory dataset, but we also analysed Functional Connectivity datasets, which comprise 60 repetitions of the same movie (30 in each block). **f** Significant pairwise and triple-wise interactions (blue dots) after applying a z-score threshold (|*z*| > 4, Methods) are shown for V1 units from an example mouse. In this case, 7.2% of the interactions were significant. The grey dots show all the interactions (including insignificant values), and the marginal distributions of significant values (blue distributions) are shown on the top (pairwise) and left (triple-wise). We calculated significant higher-order interactions from animals with at least seven recorded units and included up to 100 units. **g** Percentage of significant interactions calculated from V1 units in all mice in Brain Observatory (31 mice) and Functional Connectivity (26 mice) datasets. The average significant interactions across animals are shown in red (2.7% in Brain observatory and 5.3% in Functional Connectivity). The box plots indicate median (middle line), 25th, 75th percentile (box) and minimum and maximum (whiskers).

A limiting factor in calculating the significant HOIs is the multiple permutations of spike trains to obtain the shuffled distribution for each triplet. For large numbers of neurons, this makes the calculations extremely costly (Suppl. Fig. 1). Therefore, we developed an analytical method that calculates the mean and standard deviation of pairwise and triple-wise interactions for shuffled spike trains, given the total number of bins and the neurons’ firing rates (Fig. 1c and d, Methods). In addition to becoming computationally efficient, the analytical method has the advantage of being more accurate: numerical simulations with 100 shuffles show variation in the mean and standard deviation, whereas the analytical results match perfectly well with a large number of shuffles (i.e. 10^6^). Using our analytical method, we can obtain results equivalent to numerical simulations with arbitrarily large (infinite) numbers of shuffles of spike trains. Figure 1d, right, also shows a scaling relation between the standard deviation of θ_12_ (or θ_123_) and the number of bins (*N*), given by 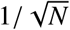. Our analytical method provides a fast and efficient method to calculate statistically significant HOIs, reducing computational time while increasing accuracy (Suppl. Fig. 1). We utilised our analytical method to calculate significant HOIs in various datasets. We first calculated HOIs for a large-scale spiking activity dataset obtained by the Allen Brain Observatory ^52^. We analysed responses to natural movies in Visual Coding Neuropixels datasets (Brain Observatory and Functional Connectivity) ^52^, Fig. 1e. The datasets comprise 20 and 60 repetitions, respectively, of the same 30-second natural movie across two presentation blocks. The spiking activities of the recorded units in V1 in response to all repetitions are shown in Fig. 1e for a sample mouse. We computed pairwise (Eq. 2) and triple-wise interactions (Eq. 4) among all combinations of three neurons. The distributions of all calculated HOIs and statistically significant ones are shown in (Fig. 1f; gray and blue dots, respectively).

Our results show that significant HOIs can be estimated from the spiking activity with an experimentally-tractable recording length (recording durations were 30 minutes for Functional Connectivity and 10 minutes for the Brain Observatory datasets). We calculated the percentage of significant HOIs out of all possible triplets for V1 units recorded in each animal (Fig. 1g). The average percentage of significant HOIs in V1 across all mice was 2.7% in Brain Observatory. This value increased to 5.3% for Functional Connectivity dataset, which contained longer recording sessions (30 minutes) due to more repetitions (60 times) of natural movies (Fig. 1f).

The HOIs obtained above were calculated for animals with at least seven simultaneous recorded units (more than 30 significant points, Methods). For animals with more than 100 recorded units, we limited the analysis to 100 units. This yielded 161,700 possible triplet combinations (choosing 3 out of 100) per animal. For larger populations, the number of combinations would be computationally prohibitive. It would be therefore useful to know how many neurons are needed to obtain a reliable estimate of the average HOI in the population. To assess that, we calculated the average HOI from sub-samples of all the neurons (Suppl. Fig. 2). The average value of HOIs remained similar when estimating it from subsamples of 20 to 100 neurons, but the variance of this estimate was reduced by increasing the subsample size. Our results suggest that the average HOI can be reliably estimated from as low as 50 neurons (Suppl. Fig. 2). Taken together, our results suggest that using CHOIR, HOIs can be reliably estimated from simultaneous recording of spiking neurons with experimentally-tractable population sizes and recording times.

### Patterns of higher-order interactions across animals and visual areas

We next asked how the estimate of HOIs change across animals and visual regions (Fig. 2a). First, we calculated significant HOIs obtained from V1 units across different mice and plotted the average significant pairwise and triple-wise interactions (Fig. 2b). To ensure a fair comparison across animals with different numbers of recorded units, we included HOIs datasets with at least 30 significant interactions in our analysis. In all the animals, we consistently observed average positive pairwise and negative triple-wise interactions for both Functional Connectivity (top) and Brain Observatory (bottom) datasets, with similar magnitudes across datasets (Fig. 2b). There was no evident relationship between average HOIs and the average firing rates, which remained relatively similar across animals (Fig. 2b, right). Our results therefore corroborate that the estimates of HOIs are reliable and consistent across animals, independent of the average firing rates.

**Fig 2.**
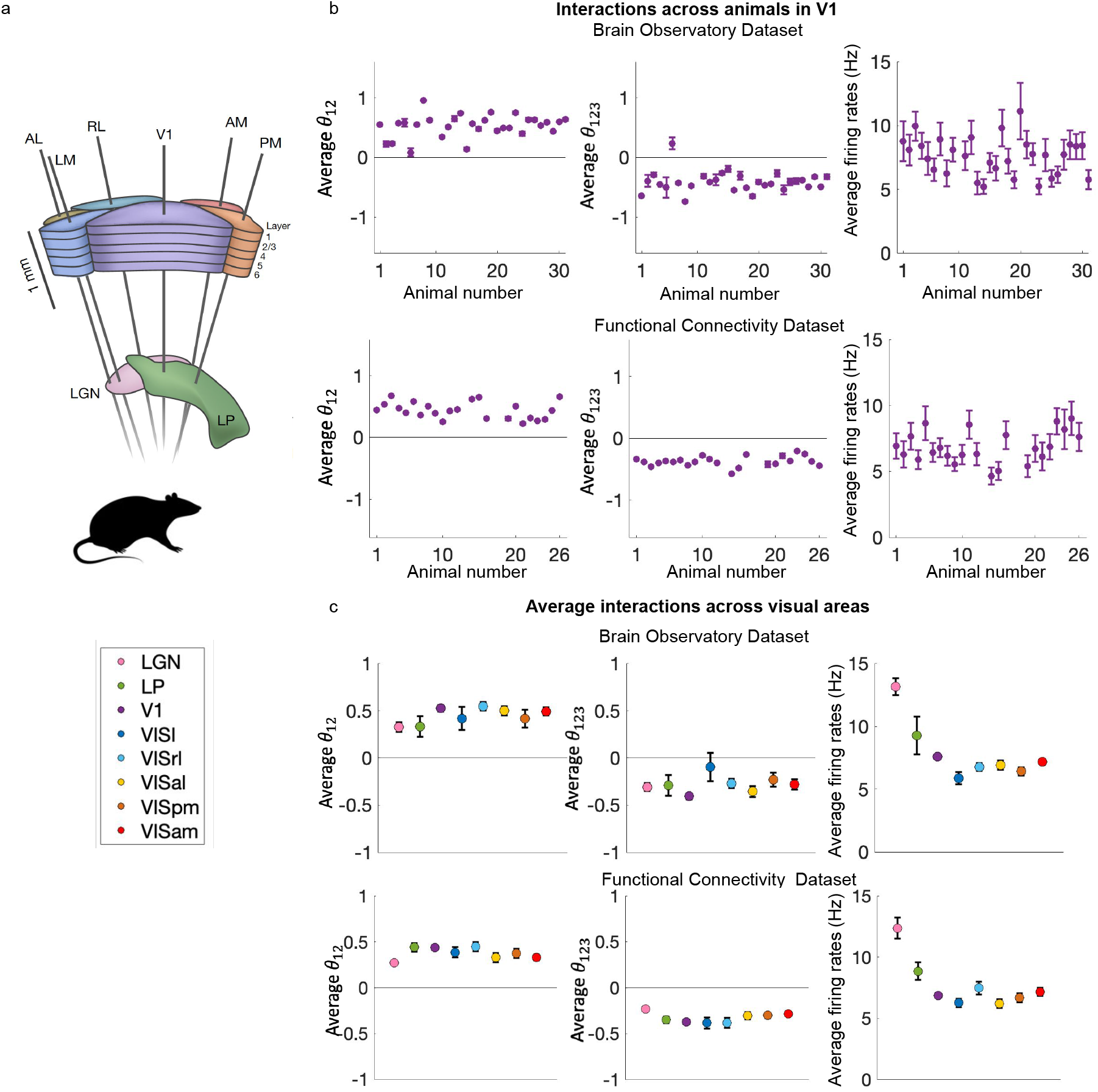
Pairwise and triple-wise interactions across animals and brain regions. **a** Schematic view of thalamus (lateral geniculate nucleus; LGN, and lateral posterior nucleus; LP), primary visual cortex (V1), and higher visual areas (lateromedial area; VISl, rostrolateral area; VISrl, anterolateral area; VISal, posteromedial area; VISpm and anteromedial area; VISam) **b** Significant average pairwise (left) and triple-wise (middle) interactions for each individual mouse in V1 for Brain Observatory (top) and Functional Connectivity (bottom) datasets (Δ = 20 ms, *p* < 0.01 by permutation test). Firing rates of neurons participating in significant interactions (right).**c** Pairwise and triple-wise interactions averaged across all mice for each region along the visual hierarchy from LGN to VISam for Brain Observatory (top) and Functional Connectivity (bottom) datasets. Firing rates of neurons participating in significant interactions for each region (right). Error bars represent ±SEM.

We also calculated the average pairwise and triple-wise interactions across visual regions (Fig 2c). The average pairwise and triple-wise interactions (across all mice) consistently showed positive and negative values respectively, across all regions in two datasets (Fig. 2c, Suppl. Fig. 3 and Fig. 4). These results were not dependent on the average firing rates of the regions (Fig. 2c, right). Moreover, they were robust to the choice of significance threshold to assess statistically significant HOIs (data not shown). Our results therefore suggest that a reliable and consistent pattern of average HOIs can be inferred across animals and brain regions.

### Higher-order interactions can reveal the nature of shared inputs

Our results so far established that reliable and significant higher-order interactions can be inferred from spiking activity. We next sought to study how HOIs can inform us about connectivity motifs underlying the neuronal interactions. Recent work showed that an analytical guide map of neural interactions (inferred from pairwise and triple-wise interactions) can reveal hidden shared input motifs ^50^. It suggests that the two-dimensional space of pairwise and triple-wise interactions provides a more refined distinction between possible motifs (Fig. 3a; as illustrated by the shaded regions and their corresponding motifs). For instance, a motif in which all three neurons receive the same shared excitatory input (excitatory-to-trio, grey shading) would be clearly distinguished from a motif in which similar excitatory inputs are shared by each pair of neurons in the triplet (excitatory-to-pairs, magenta) (Fig. 3a).

We projected the average significant pairwise and triple-wise interactions of V1 neurons onto the guide map for each animal (Fig. 3b). For most of the animals, the data points clustered in the area corresponding to the motif of excitatory inputs to each pair within the triplet (Fig. 3b). The distribution of data points had minimal overlap with regions belonging to alternative motifs, and the data was far away from the control analysis obtained from shuffled spike trains, which clustered close to zero (Fig. 3b, green dots). This result was similar for Brain Observatory dataset (Suppl. Fig. 5a). We also repeated our analysis for a smaller time window of spiking analysis (Δ = 10 ms) and obtained similar results (Suppl. Fig. 6). Our results, therefore, suggest that CHOIR can be used to distinguish reliable and specific motifs of neural activity.

**Fig 3.**
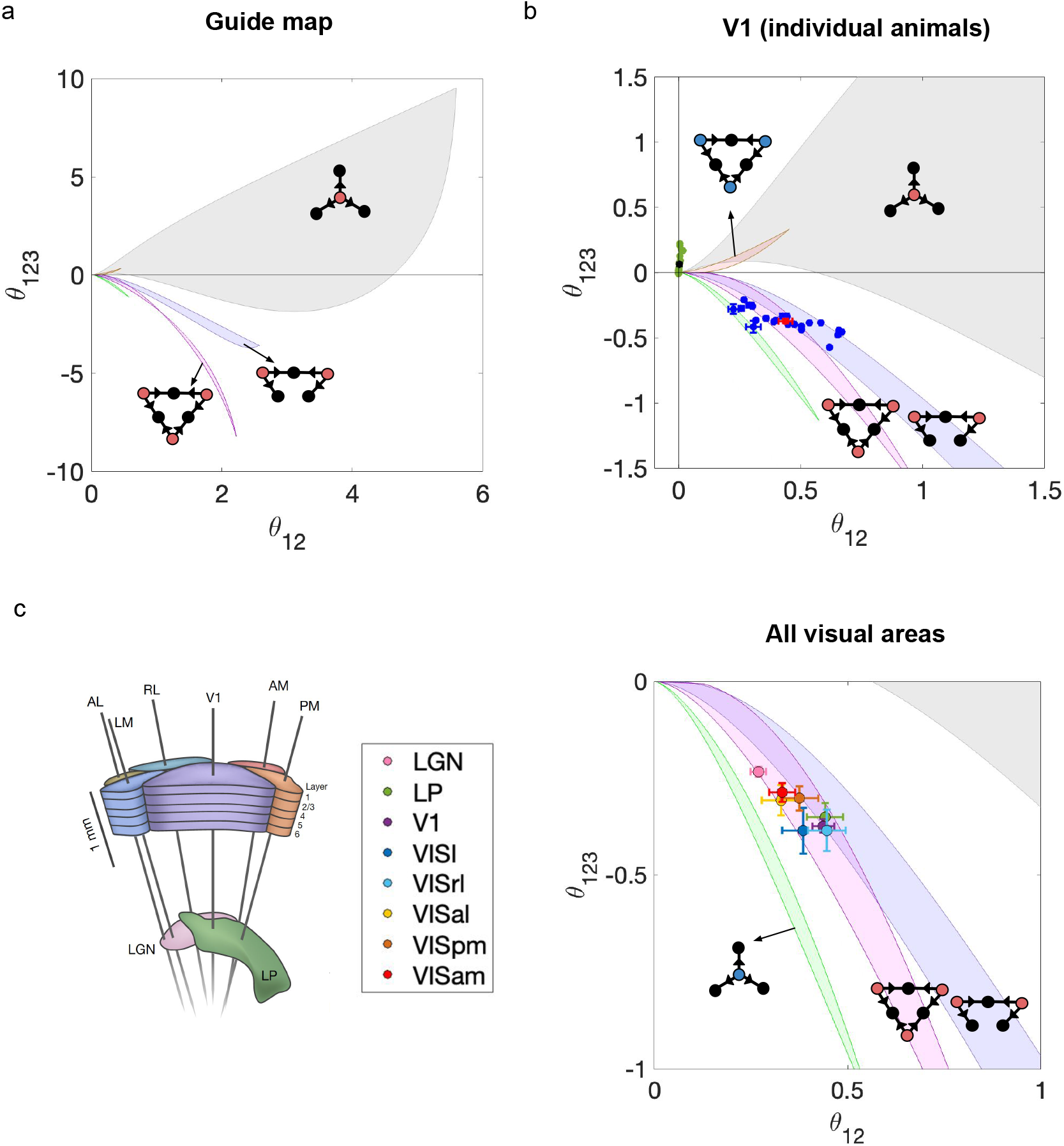
Analytical guide map of neural interactions to reveal underlying motifs. **a** Guide map for revealing hidden shared inputs in the plane of pairwise and triple-wise interactions ^50^. Each colored region is associated with a specific motif with hidden shared inputs. The grey area corresponds to excitatory shared inputs given to three postsynaptic neurons. The pink and blue areas define regions for excitatory shared inputs to pairs of neurons out of three (symmetric and asymmetric). **b** The interactions of each individual mouse V1 data (blue) and the average over all mice (red) on the guide map of neural interactions. The green dots are interactions from shuffled spike trains and the black dot shows the mean ±SEM. **c** Average pairwise and triple-wise interactions across thalamus and visual areas (from LGN to VISam, Functional Connectivity dataset) plotted on the guide map. The time window is Δ = 20 ms and error bars represent SEM (*p* < 0.01 by permutation test).

Next, we analysed the pattern of HOIs distributions across visual regions. We repeated the across animal analysis for other regions and found that most of the data points were again located in the region corresponding to the excitatory-to-pairs motif, although there were different levels of scatter across animals in different regions (Suppl. Fig. 7). We calculated the average HOIs values across animals and plotted them for each region on the HOIs guide map. The data consistently fell within the region attributed to the symmetric excitatory-to-pairs motif, notably, this represents a very narrow area in the interaction plane (Fig. 3c). The result was the same across the Brain Observatory dataset (Suppl. Fig. 5b and Suppl. Fig. 8). The dominant inferred motif across brain regions is thus consistent with our previous analysis across individual animals in V1 (Fig. 3b), and the results recently reported in monkey and mouse V1^50^ too. Our analysis therefore suggests that HOIs can be used to estimate the underlying motifs of neural interactions, and that the excitatory-to-pairs motif emerges as the basic prevailing motif across visual areas, individual animals, and potentially species.

### Higher-order interactions can differentiate between behavioural states

The dynamics of neural circuits are strongly modulated by the behavioural states of animals ^57–61^, affecting visual coding across multiple brain regions ^62^ and contributing to representational drift ^61^. These behavioural state-dependent changes can in turn alter the interactions among neurons. To examine how changes in behavioural states affect neural interactions, we analysed HOIs in different behavioural states. We categorised different repetitions of the natural movie as belonging to running or stationary states (Methods), based on the animal’s running speed. We calculated significant HOIs from responses to natural movies during stationary and running states separately. The result revealed a clear change in pairwise and triple-wise interactions between behavioural states in V1 (Fig. 4b) and in all visual areas (Fig. 4c). Animals exhibited predominantly stronger positive pairwise and negative triple-wise interactions in stationary trials compared to running trials (Fig. 4b, data from V1). This pattern persisted when averaging across mice for each region (Fig. 4c). The same results are also observed in another dataset (Brain Observatory, Suppl. Fig. 9 a, b). To gain a better understanding of the changes between running and stationary states, we went beyond average values of pairwise and triplet-wise interactions and investigated how the distribution of interactions changes. To this end, we analysed the distribution of interactions across all four possible quadrants of positive (+) or negative (-) pairwise (p) and triple-wise (t) interactions (1: p+,t+; 2: p+,t− ; 3: p−,t+; 4: p−,t−) (Fig. 4d,e). This analysis revealed that running data appeared in all four quadrants (Fig. 4d, left), while stationary data clustered primarily in the quadrant corresponding to positive pairwise (p+) and negative triple-wise (t-) interactions. This pattern held for V1 and when averaging across all visual areas (Fig. 4d,e, left). The same result was also obtained for the Brain Observatory dataset (Suppl. Fig. 9). The presence of negative pairwise interactions in the running state contributes to the overall reduction in the average pairwise interaction strength (Fig. 4d,e, right). These negative pairwise interactions can arise from circuits with recurrent inhibition (more specifically, lateral inhibition), which induce *P*_10_ and/or *P*_01_ patterns and generate negative pairwise interactions (Eq. 2). Analysis of pairwise and triple-wise interactions for stationary and running in another bin width (Δ = 50 ms) shows that the result persists across different time windows (Suppl. Fig. 13, and Suppl. Fig. 17). Analysis of higher-order interactions can therefore differentiate between stationary and running states in animals, helping to reveal their corresponding circuit motifs.

**Fig 4.**
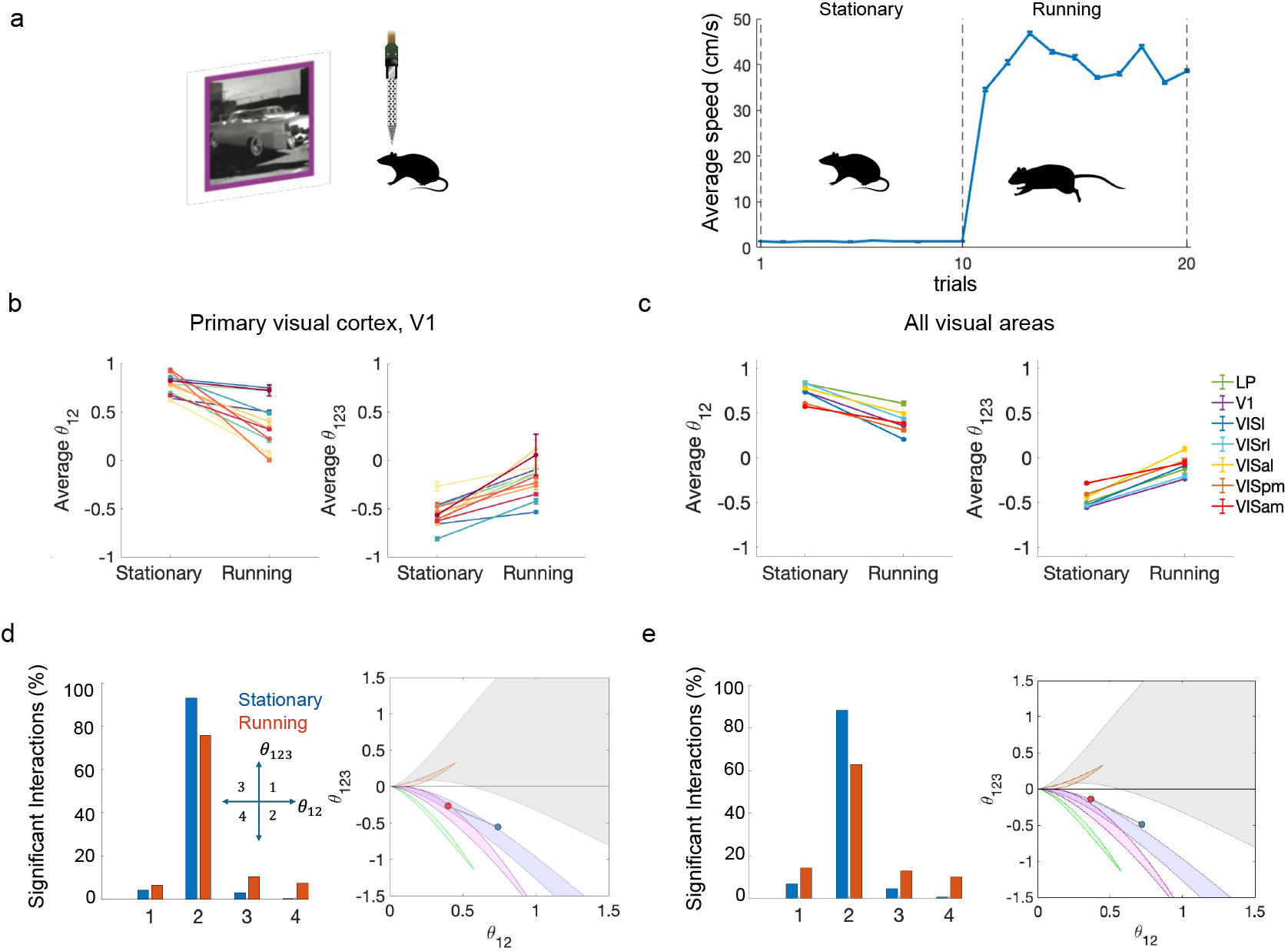
Higher-order interactions differ in stationary and running states. **a** Left: Schematic of Neuropixels recordings from mouse visual cortex during natural movie presentation (Allen Brain Observatory, Functional Connectivity dataset). Right: The average speed for each trial is shown for a mouse. Trials with speed less than 2 cm/s were considered stationary, and the rest as running. **b** Average pairwise (left) and triple-wise (right) interactions for stationary and running trials of each mouse in primary visual cortex (V1). Between stationary and running states, pairwise and triple-wise interactions differ significantly in all thirteen animals (*p* < 0.01 by permutation test), while firing rates show no significant difference across six animals (Suppl. Fig. 10). **c** Pairwise (left) and triple-wise (right) interactions averaged over mice across visual hierarchies. Except for pairwise interactions in VISl and VISrl, all other interactions differed significantly between stationary and running states. Firing rates of neurons for each region are shown in Supplementary Fig. 10. Error bars in **b** and **c** represent SEM. The results of each animal for other visual regions are shown in Supplementary Fig. 11. **d** Left: percentage of significant interactions for stationary and running states in each quadrant of triple-wise versus pairwise plane in V1. Right: Average pairwise and triple-wise interactions across all animals from panel b (V1) for stationary (blue) and running (red) states, plotted on the guide map. **e** Left: Percentage of significant interactions for stationary and running states in each quadrant of the interaction plane (triple-wise versus pairwise plane), averaged across all visual areas. The percentage for each visual area across hierarchy is shown in Supplementary Fig. 16. Right: Average pairwise and triple-wise interactions across all visual areas from panel c for stationary (blue) and running (red) states, plotted on the guide map.

### Balanced networks with structured connectivity replicate HOI patterns in stationary and running states

The balance of excitation-inhibition in networks plays a crucial role in shaping network dynamics and assembly formation ^63–67^. To identify the network properties that can generate patterns consistent with the observed pattern of HOIs in the data, we simulated large-scale balanced spiking networks (Fig.5, Methods). Our networks consisted of 1000 leaky integrate-and-fire (LIF) neurons with 800 excitatory and 200 inhibitory neurons ^64,68,69^. Neurons were randomly connected with probability *p*_*c*_ = 0.1. The network parameters were set in a way to replicate the asynchronous-irregular (AI) regimes of cortical activity with irregular firing (Methods) ^64^. We simulated the spiking activity of networks for 100 seconds and analysed HOIs in the same way as performed for the experimental data. We found that typical balanced random networks in the AI regime failed to reproduce the experimentally observed HOI patterns (suppl Fig. 20). We then systematically varied the degree of recurrent excitation and inhibition in the networks and calculated HOIs. We specifically tested whether increasing recurrent inhibition may generate the expected patterns. However, the empirical patterns of HOIs were not observed in networks with any combination of excitation-inhibition balance (Suppl. Fig. 18) nor by increasing the connectivity strength under biologically plausible firing rate regimes (Suppl. Figs. 19-20).

**Fig 5.**
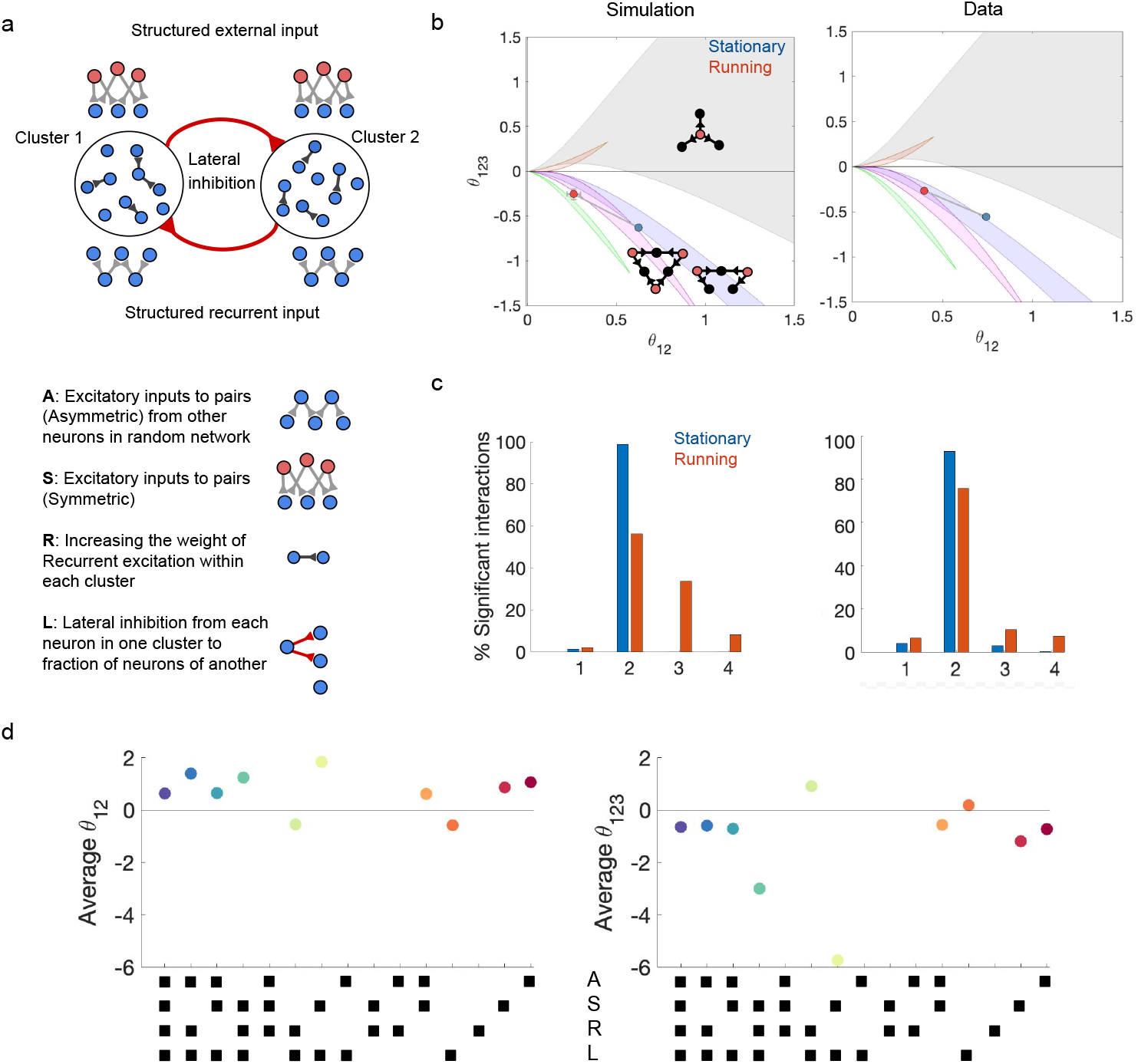
The suggested circuit motifs for stationary and running. **a** Random network simulation of 800 excitatory and 200 inhibitory spiking neurons with sparse connectivity (*p*_*C*_ = 0.1) in the asynchronous irregular regime. Two clusters of 40 neurons have been created, by enhancing their within-cluster recurrent excitation (*w*_*E*_ = 5) and cross-cluster inhibition (with probability *c* = 0.4 and weight *w*_*L*_). Structured excitatory inputs to pairs within each triplet are provided through external (symmetric, red) and recurrent (asymmetric, blue) pathways (Poisson rate λ = 5*Hz* and *w*_*A*_ = 10). **b** The network replicates stationary result when the lateral inhibitory weight is set to *w*_*L*_ = 5 (left, blue dot), similar to stationary result in data (right, blue dot). By increasing this weight to *w*_*L*_ = 15, the network produces results that resemble running state (red dot). **c** Significant interactions for stationary and running states in each quadrant of the triple-wise versus pairwise plane for simulation (left) and V1 data (right). **d** Network elements’ contribution to neuronal pairwise (left) and triple-wise (right) interactions. The full model consists of all four elements: (**A**) Recurrent excitatory-to-pairs (asymmetric) within triplets of each cluster from other neurons in the network. (**S**) Excitatory inputs to pairs (symmetric) within triplet of each cluster. (**R**) Increasing the weight of recurrent excitatory connections within each cluster. (**L**) Lateral inhibition from each neuron in one cluster to a portion of neurons in another. Other parameters are: *V*_θ_ = 20 *mV, V*_*reset*_ = 10 *mV, V*_*rest*_ = 0 *mV*, τ_*m*_ = 20 *ms*, τ_*delay*_ = 1.5 *ms*, τ_*re−f*_ = 2 *ms, w*_*e*_ = 0.1, *w*_*i*_ = −0.5, *g* = 5 and Δ = 20 *ms*.

Next, we analysed HOIs in balanced networks equipped with structured connectivity ^70–73^. We simulated networks with clustered connectivity patterns, where (1) recurrent connections were organised according to a Mexican hat profile with stronger recurrent excitatory connectivity within each cluster and stronger lateral inhibition between them, and (2) excitatory inputs given to pairs of neurons in each cluster (Methods; Fig.5a). Interestingly, the structured networks revealed HOI patterns similar to the experimental results (Fig.5b). The stationary data showed that the average pairwise and triple-wise interactions were located in quadrant 2 (Fig.5b, right, blue circle). During running, the average pairwise shifted leftward and the average triple-wise moved upward (Fig.5b, right, red circle) relative to the stationary data. Our simulated networks could, therefore, replicate both the difference between the stationary and running states and the direction of these changes: weak lateral inhibition reproduced the HOIs patterns observed in the stationary state, whereas stronger lateral inhibitory weights replicated the HOI patterns characteristic of the running state (Fig.5b and c).

To elucidate how each component of the structured networks contributes to HOI patterns, we simulated networks with different combinations of connectivity elements. Specifically, we simulated random networks and added the following four components: excitatory input to pairs, consisting of symmetric excitatory to pairs(S) as external input, and asymmetric excitatory to pairs(A) input from a fraction of other neurons in the network; stronger weights of existing recurrent excitation within clusters (R); and lateral inhibition (more than random) across clusters (L) (Methods). We simulated networks with all 15 combinations of parameters, calculated pairwise and triple-wise interactions in each of them, and compared the results with the full network where all elements were present (S=1, A=1, R=1, L=1) (Figure 5 d). Among all four network elements, excitatory inputs to pairs motifs (S and A, Figure 5a, bottom) led to generally positive pairwise and negative triple-wise interactions. This excitatory-to-pairs motif was also revealed as the dominant motif from data analysis (Fig.3 c). However, when existing random recurrent excitation was strengthened (R) in random networks containing these motifs (S and A), no significant HOIs were detected, suggesting that strong excitatory recurrent connections disrupted the structured common input effects (excitatory to pairs) responsible for coordinated firing patterns.

The presence of lateral inhibition (L) alone in the network generated negative pairwise and positive triple-wise interactions. However, by adding this motif (L) to symmetric excitatory inputs to pairs (S), the network produced the strongest positive pairwise and negative triple-wise interactions. We therefore conclude that, to replicate the results of interactions consistent with empirical data for stationary and running, all four motifs (S, A, R, and L) need to be combined. Interestingly, all connectivity motifs have been reported in cortical networks ^2,70–72,74–79^. In such networks, increasing the weight of lateral inhibition replicates the change in HOIs consistent with transitions from stationary to running states (Fig. 5b). Our results therefore suggest that the dynamic nature of neural interactions and their modulation by the brain state can be recapitulated in large-scale balanced networks, when they are equipped with structured recurrent and feedforward connectivity motifs.

### Higher-order interactions can differentiate distinct neural ensembles

Our analysis before suggested that shared excitatory inputs to pairs and lateral inhibition between them were important network motifs to explain the pattern of HOIs inferred from Allen Brain datasets. However, this inference was indirect, as such ensembles and their lateral inhibition were not directly probed in the data. To verify our method directly, and to go beyond one dataset, we analysed HOIs in a recent dataset, which specifically studied ensembles of neurons in the visual cortex ^54^. In this dataset each stimulus (drifting grating with a given direction) modulated the activity of a subset of V1 neurons (“ensembles”), while the remaining neurons showed no change in activity (Fig 6a, non-participants, grey circles). Within each ensemble, some neurons increased their activity during the stimulus (called “onsemble”, blue circles), while others decreased their activity (called “offsemble”, red circles). We therefore asked whether we can distinguish between these ensembles by analysing their HOI patterns, and whether the hypothesised lateral inhibition can be detected.

**Fig 6.**
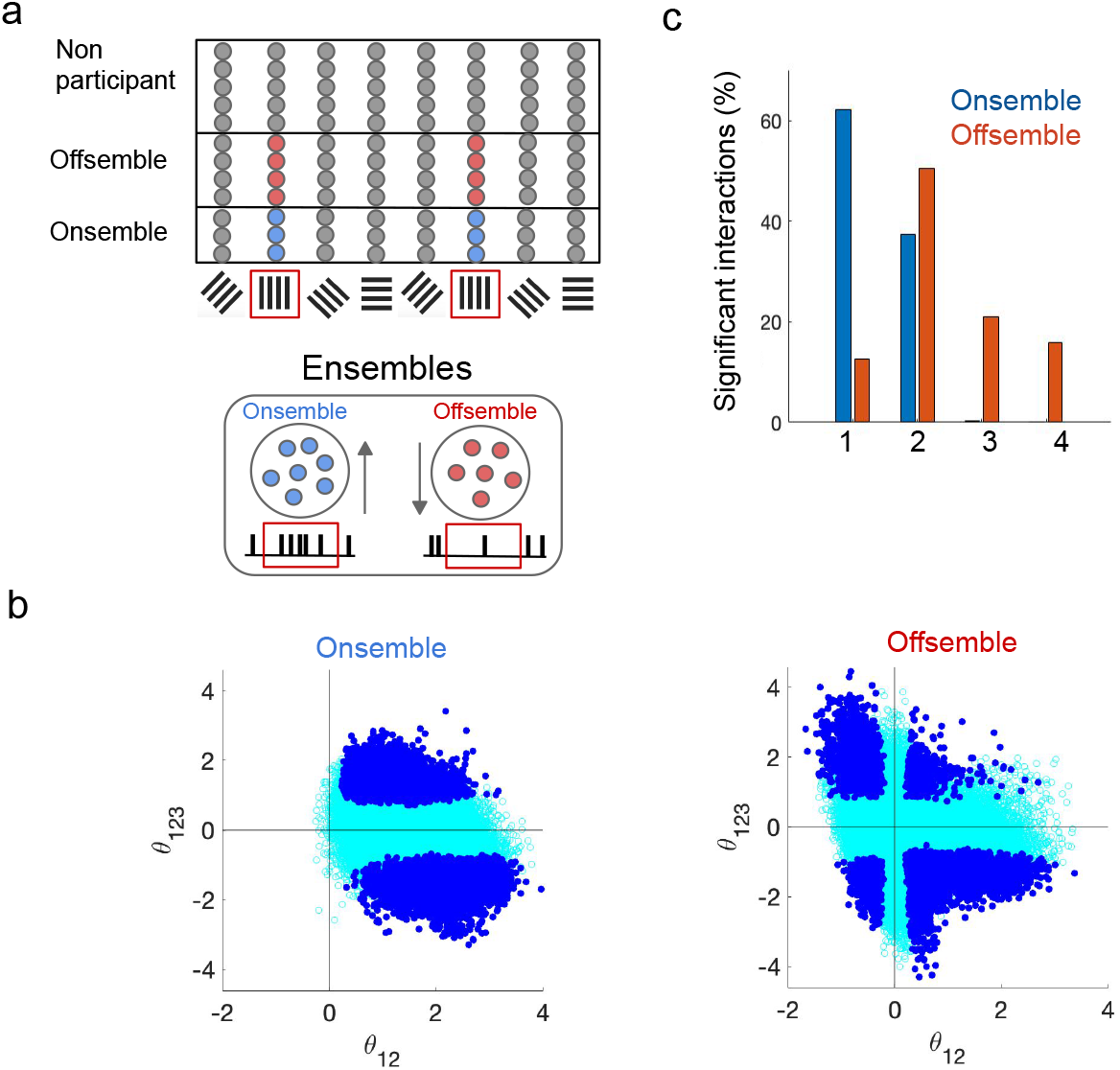
Pairwise and triple-wise interactions can distinguish different ensembles of neurons. **a** Schematic of ensembles of neurons which get activated (onsemble) or deactivated (offsemble) during preferred stimulus presentation. **b** Significant pairwise and triple-wise interactions (blue) for offsemble and onsemble neurons in one ensemble of a mouse in response to drifting grating stimuli (*p* < 0.01 by permutation test). Data from Pérez-Ortega et al. ^54^.**c** Significant interactions across four quadrants in the triple-wise versus pairwise interaction plane for onsemble and offsemble neurons, aggregated from twelve mice, each with five ensembles.

Significant interactions were calculated for each onsemble and offsemble groups separately. For each mouse, we examined five distinct neuronal ensembles of each type. We observed that offsemble neurons exhibited both negative and positive pairwise interactions (Fig. 6b, bottom panel). In contrast, onsemble neurons showed exclusively positive pairwise interactions (Fig. 6b, top panel). Using data aggregated from all twelve mice, we also analysed the distribution of statistically significant interactions (Suppl. Fig. 21) in the four quadrants of pairwise versus triplet-wise interaction plane (Fig. 6c). The majority of significant interactions for onsemble neurons were in quadrants with positive pairwise interactions (i.e. quadrants one and two). However, for offsemble neurons, considerable fractions of data were in quadrants with negative pairwise interactions (quadrants three and four), and also the fraction of data in positive pairwise and triple-wise interactions (quadrant one) is decreased compared to onsemble neurons. Our analysis showed that offsemble neurons, in contrast to onsemble neurons, exhibit negative pairwise interactions, revealing lateral inhibitory mechanisms. The experimental results suggested that offsemble neurons might have some inhibitory mechanisms due to a faster decrease in calcium transients compared to onsemble neurons ^54^. This observation aligns with our finding of negative pairwise interactions among offsemble neurons. Therefore, our analysis of neuronal interactions among offsemble neurons suggests that lateral inhibition can be the inhibitory mechanism operating within this population while being absent among onsemble neurons.

The majority of onsemble neuronal interactions are located in quadrant one (positive pairwise and positive triple-wise interactions), where excitatory-to-trio motifs can explain this pattern (according to our motif guide map, Fig.3a). In contrast, offsemble neurons show considerable decreased excitatory-to-trio motif and instead exhibit increased lateral inhibition. This difference in network motifs between onsemble and offsemble populations can suggest a mechanistically distinct circuit organisation depending on the functional class, with onsemble neurons supporting cooperative amplification while offsemble neurons implementing competitive suppression among neurons.

### Higher-order interactions can distinguish sleep and awake states

We further applied CHOIR to an additional dataset to assess whether significant HOIs may differentiate between sleep and wakefulness, two brain states with dramatically different network dynamics ^80–82^. We analysed sleep and wakefulness data from mouse V1^53^, where neural spiking activities were recorded simultaneously during both states. We focused on non-REM (N-REM) sleep, which provided longer continuous recording sessions than REM sleep. By calculating significant pairwise and higher-order interactions across all neuron triplets, we found that the N-REM sleep state shows stronger positive pairwise and negative triple-wise interactions than awake state in most animals. Therefore, we could effectively distinguish the two behavioural states (Fig.7a).

**Fig 7.**
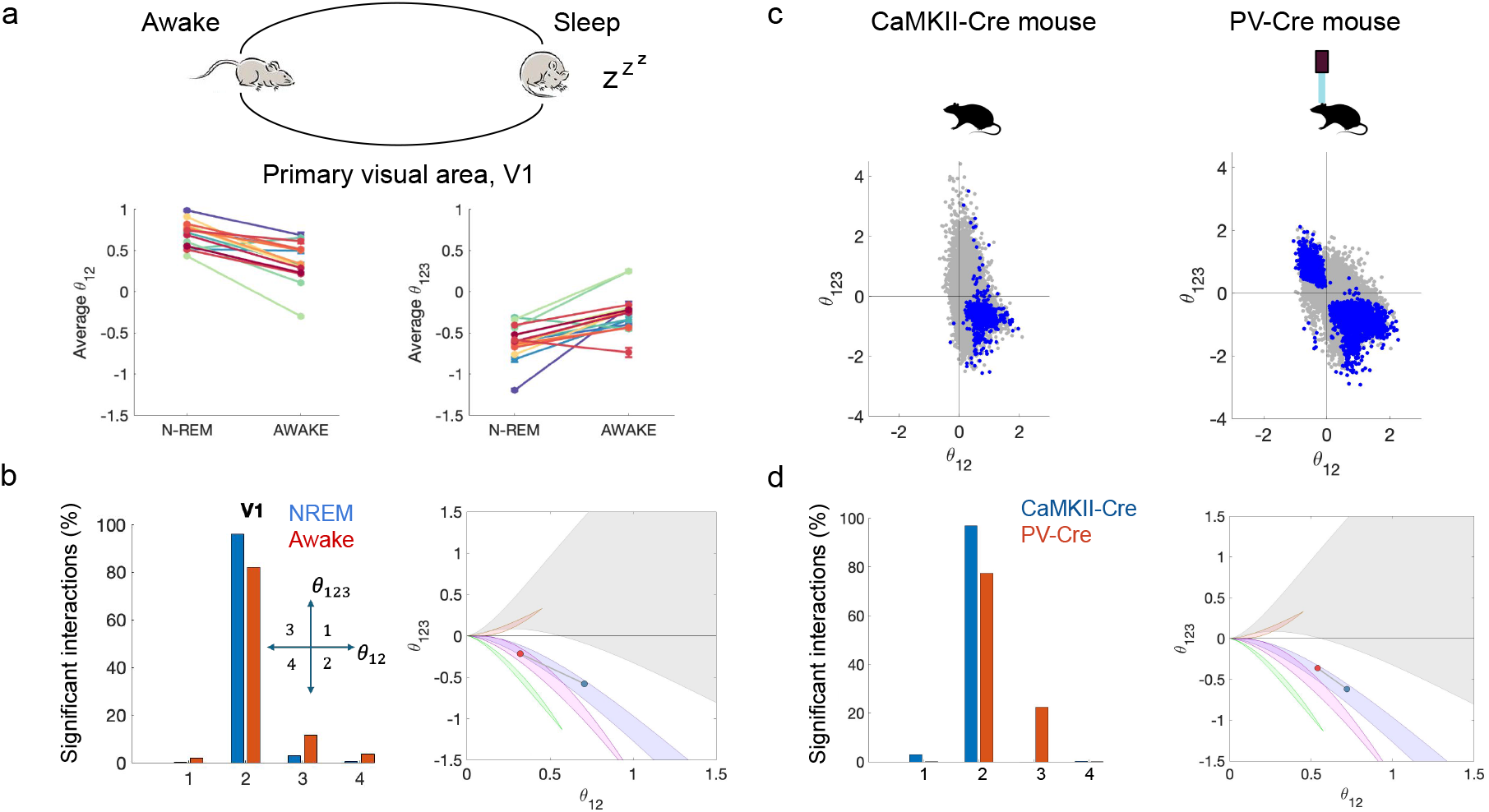
Pairwise and triple-wise interactions can distinguish states and ensembles of neurons. **a** Sleep (N-REM) and awake data ^53^ can be differentiated by pairwise and triple-wise interactions (Top). **b** Significant interactions for sleep and awake states in each quadrant of triple-wise versus pairwise plane (left). Average pairwise and triple-wise interactions across all (17) animals in V1 for sleep (N-REM, blue) and awake (red) states, plotted on the guide map (right). **c** Significant interactions in two mice in sleep (N-REM) state (blue dots). In one mouse, Parvalbumin-expressing inhibitory inter-neurons (PV-Cre, right) are activated by optogenetic stimulation whereas the other has no stimulation (left). **d** Percentage of significant interactions in each quadrant (left) for optogenetical stimulation of PV neurons and control. Average pairwise and triple-wise interactions for PV-Cre (blue) and control mice (red). The bin size is Δ = 20 *ms* (*p* < 0.01 by permutation test).

We then asked what the state-dependent differences in interactions suggest about the dynamical motifs that have the strongest influence on network dynamics during each state. To this end, we examined how statistically significant interactions were distributed across the four quadrants of the pairwise versus triplet-wise interaction plane. The distribution of the interactions revealed clear differences between the two states (Fig.7b). During wakefulness, interactions were scattered across multiple quadrants, including quadrants three and four (characterised by negative pairwise interactions) alongside quadrant two (Fig.7b, left). In contrast, interactions during N-REM sleep were concentrated primarily in quadrant two (i.e., positive pairwise and negative triplet-wise interactions), a pattern explained by the excitatory-to-pairs motif.(Fig.7b, blue dot). During wakefulness, lateral inhibition is strengthened, shifting the distribution of average interactions leftward from the sleep state to the awake state (Fig.7b, blue and red dots). These results demonstrate that analysing HOIs can distinguish between Non-REM and awake states, similarly to the distinction between stationary and running states.

In the same dataset ^53^, we also examined the interactions by comparing a mouse with optogenetically activated Parvalbumin-expressing interneurons (PV neurons) with a control animal during sleep (Fig.7c). Optogenetic activation of PV neurons using blue light in PV-Cre mice produced predominantly negative pairwise interactions, compared to the control condition. This finding directly demonstrates that enhanced lateral inhibition, driven by active PV neuron recruitment, generates stronger negative pairwise interactions relative to control (Fig.7d). Taken together, our analysis across behavioural states suggests that HOI analysis can distinguish between different states and reveal circuit motifs, such as lateral inhibition, that are preferentially active in specific states or ensembles. Optogenetic stimulation of PV neurons confirmed that negative pairwise interactions serve as a signature of this inhibitory motif. These findings illustrate how HOI analysis can uncover specific circuit mechanisms that vary with behavioural states at the neuronal population level.

## Discussion

We introduced CHOIR (Circuit motifs from Higher-Order Interactions in neural Recordings), a method that overcomes fundamental bottlenecks in analysing higher-order interactions from large-scale neural recordings: the statistical challenge of reliably estimating significant HOIs from limited experimental data, and the computational expense of calculating interactions across all neuron combinations. By assessing statistical robustness from random permutations and replacing numerical shuffling with an exact analytical approach, CHOIR reduces computational costs by orders of magnitude while ensuring reliable detection of significant HOIs. Applying CHOIR across multiple datasets, we demonstrated that HOIs reliably distinguish between brain states, reveal functional circuit motifs, and identify neuronal assemblies. By linking HOIs to the underlying connectivity, we showed how circuit architecture supports state-dependent dynamics, and this was validated through analysis of causal manipulation experiments.

Our systematic analysis of pairwise and higher-order interactions in mouse visual cortex revealed consistent organisational principles governing neuronal circuit function. Across all visual areas, we observed a striking pattern: positive pairwise interactions coupled with negative triple-wise interactions. This signature, when interpreted through the guide map, points to a dominant circuit motif in which excitatory inputs are given to pairs of neurons within triplets. Notably, this organisational principle is not unique to rodents; previous work on macaque V1 demonstrated an identical motif ^50^, and anatomical studies of V1 and somatosensory cortex similarly report over-representation of this motif ^1,2^. These converging lines of evidence from multiple species and methodological approaches suggest that excitatory shared inputs to pairs represents a fundamental building block of cortical circuits.

A key finding from our analysis is that the strength and sign of neural interactions dynamically shift between behavioural states. By comparing average interactions during stationary and running ^52^, sleep and awake ^53^, and different ensembles of neurons ^54^, we identified distinct signatures and revealed the influential motifs for each state (Fig.3, Fig.6, and Fig.7). This behavioural sensitivity reveals the computational flexibility of cortical circuits: rather than operating as fixed assemblies, neural populations actively reconfigure their interaction patterns in response to behavioural states. We observed the emergence of negative pairwise interactions for running versus stationary, awake versus sleep states, and also among “offsemble” neurons in contrast to onsembles. We interpret this signature as evidence for lateral inhibition operating between neurons. This contrasts with other inhibitory mechanisms that would simultaneously suppress multiple neurons (blanket inhibition), which instead generate positive pairwise interactions. Our method can distinguish these mechanistically distinct inhibitory processes and suggests a mechanism for the observed interactions.

The lateral inhibitory motifs observed during locomotion align with known neuromodulatory mechanisms in cortex. Previous studies showed that VIP interneurons disinhibit parvalbumin-expressing (PV) neurons by inhibiting somatostatin-expressing (SST) neurons ^57,59,60,83^, enabling PV neurons to inhibit pyramidal neurons. This increased PV-to-pyramidal inhibition is consistent with the lateral inhibitory motifs we detected during running. On the other hand, some studies showed that SST interneurons, which are known to laterally inhibit pyramidal neurons ^74^, remain functionally engaged during running ^59,83–87^, where they stabilise dynamics of the network in concert with PV neurons ^88^. In this case, our observed lateral inhibitory signatures likely reflect the combined contributions of both PV and SST inhibitory motifs during locomotion. The inhibitory architecture during running can also involve LAMP5+ neurogliaform cells, a population that generates spontaneous activity correlated with locomotion speed ^89^. Unlike VIP interneurons that selectively target SST cells, LAMP5+ cells widely inhibit both excitatory and inhibitory populations. If this inhibition manifests as lateral suppression among excitatory neurons, it can generate the negative pairwise interactions we observed. Our results suggest that this lateral inhibitory mechanism (rather than population-level silencing) predominates during running states.

We replicated the distinct HOI signatures of stationary and running states through simulation of structured, balanced spiking networks of excitatory and inhibitory neurons. Random networks in the asynchronous-irregular regime ^64^ did not show significant triple-wise or pairwise interactions under biologically plausible firing rate regimes (Suppl. Fig. 18-20). Other balanced networks with different dynamical properties may in principle generate such interactions ^63,90^, but the firing rates in these chaotic networks are typically high and outside the biologically plausible range for mouse visual cortex (1 −10 Hz, Allen data) ^57^. By adding structured excitatory input to pairs within each cluster and lateral inhibition between clusters, we found that the empirical results can be replicated in biological ranges of firing rates. The same network mechanism can describe the observations in non-REM sleep and awake data: clusters receiving structured inputs within themselves (i.e., excitatory-to-pairs inputs) can replicate the result of sleep, while for awake state adding strong lateral inhibition between the clusters can generate the observation. Consistent with our findings, recent experiments on multiple scales (from macroscale recordings (fMRI, EEG) ^81,82,91^to cellular-level measurements ^80^) demonstrate that brain networks are more segregated into modules during sleep and anaesthesia, but more integrated during wakefulness ^80^. This convergence of empirical findings, circuit-level analysis, and computational modelling further supports our mechanistic interpretations. We also found that optogenetic stimulation of PV neurons can induce negative pairwise and positive triple-wise interactions, while in the control case, HOIs did not show negative pairwise interactions. This causal manipulation confirms that active PV neurons can create lateral inhibitory motifs among neurons detected by our HOI analysis (Fig. 7).

A recent study examined recurrent connectivity motifs among triplets of neurons using cross-correlogram analysis ^92^. However, this approach is limited to detecting recurrent connections and cannot resolve shared inputs converging on neuron pairs and trios. In contrast, our method overcomes this limitation by linking observed interaction patterns with underlying circuit motifs through an analytical guide map. Critically, the cross-correlogram approach in this work (which identified six overexpressed motifs with all excitatory links) did not reveal any difference between stationary and running states, whereas our analysis uncovers state-dependent lateral inhibitory motifs that emerge selectively during locomotion. This discrepancy highlights the power of interaction-based analysis in revealing hidden circuit dynamics that correlate with behaviour.

The circuit motifs that we identified provide mechanistic insights into how cortical circuits are dynamically gated during behavioural transitions. Top-down feedback during locomotion can disinhibit PV neurons, which then implement lateral inhibition among excitatory populations. This feedforward-feedback organisation enables rapid switching between functional states from stationary to locomotion. Such state-dependent gating may reflect broader principles of cortical computation, whereby the same anatomical circuits support distinct functional modes depending on behavioural context and neuromodulatory drive. Understanding these state-dependent interaction signatures may also have implications for detecting and diagnosing brain diseases. Previous studies showed that HOIs can distinguish between amnesic and control mice ^40^ and can differentiate healthy patients from clinical groups (patients with different levels of consciousness) ^44^. Our method can therefore help to reveal circuit level dysfunction underlying cognitive deficits even when individual neural firing rates appear relatively preserved. For instance, in Alzheimer’s disease, tracking the change of higher-order interactions and their relevant motifs could provide early biomarkers of circuit dysfunction before widespread cell loss occurs. Our approach holds particular promise for diseases characterised by altered excitatory-inhibitory balance, such as Autism, Schizophrenia, and Parkinson’s ^93–100^, where motifs may emerge differently across behavioural states. In Parkinson’s disease, for instance, unusual patterns of synchronisation in basal ganglia circuits might be characterised by disrupted interaction motifs, potentially guiding therapeutic interventions. For epilepsy, mapping pathological interaction cascades could potentially identify critical nodes in seizure generation networks. We also showed that applying our framework to sleep-awake states can reveal how motifs reorganise from the patterns of non-REM sleep to active wakefulness. Also disrupted sleep-related motifs can mark several diseases, ranging from psychiatric conditions like depression to neurodegenerative disorders such as Alzheimer’s and Parkinson’s.

More generally, our analytic approach to obtaining statistically significant HOIs can have broad applications in computational biology and beyond. The closed-form relations we derive for the mean and variance of null distributions of pairwise and triple-wise interactions in binary sequences enable fast, scalable detection without the computational burden of permutation testing. In gene regulatory networks, these results facilitate rapid identification of co-expressed gene pairs and synergistic gene triplets, with natural extensions to genome-wide epistasis mapping and CRISPR screens. Similar benefits arise in epidemiology, where it enables systematic discovery of disease comorbidity pairs and multimorbidity triplets; and in climate science, where it supports identification of teleconnections and compound extreme events. Across all these domains, replacing expensive permutation tests with analytic null expressions makes HOIs detection both computationally tractable and statistically rigorous.

In summary, our work establishes pairwise and higher-order interaction analysis as a powerful framework to bridge the gap between large-scale neural recordings and mechanistic circuit understanding. By decomposing complex interaction patterns into interpretable motifs, we provide a principled method to extract circuit-level insights from population recordings. Extending this approach to other brain regions and behavioural states can reveal how general principles of circuit organisation scale across the brain and how they support diverse computations. Ultimately, our study demonstrates that neural coding emerges not just from the activity of individual neurons, but from the precise patterns of coordination and competition among neural ensembles. By systematically characterising these interaction patterns and linking them to identified circuit motifs, our analysis advances our understanding of how the brain’s networks underlie neural dynamics and behaviour.

## Methods

### Pairwise and triple-wise interactions

We calculated the interactions among the neuronal activities. For two neurons, the probability mass function of individual and joint activity among neurons in exponential representation is ^15,27^:

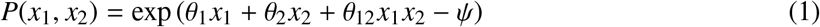

where *x*_*i*_ represents the binary activity of neuron *i*, which can be either active (*x*_*i*_ = 1) or silent (*x*_*i*_ = 0), and *ψ* is the normalization factor. The θ_*i*_ terms are canonical parameters, and θ_12_ represents the pairwise interaction between two neurons ^15,18,56^:

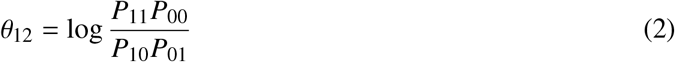

Here, *P*_11_ is the probability of both neurons being active simultaneously, *P*_10_ and *P*_01_ are the probabilities of one neuron being active while the other is silent, and *P*_00_ is the probability of both neurons being silent. Extending this formalism to three neurons, the probability mass function reads:

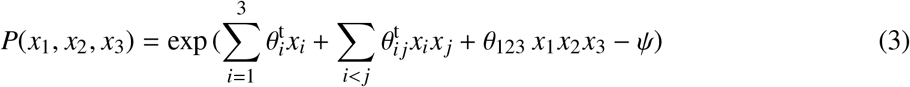

where 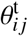 represents the pairwise interaction between two neurons in the context of three neurons, and *θ*_123_ is the triple-wise interaction between three neurons ^17,18^:

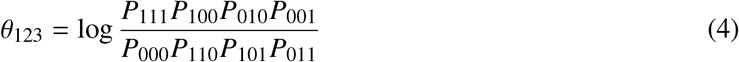

The terms *P*s represent the probabilities of pattern occurrences. For example, *P*_111_ is the probability that all three neurons are active simultaneously.

### Detecting significant interactions

Here, we provide a detailed description of how we assessed the statistical significance of pairwise and triple-wise interactions among neuronal spike trains (Fig. 1). For each region, we selected animals that had at least seven recorded units (more than 30 points), with an upper limit of 100 units. To have a better estimate of the interactions, we concatenated all the trials for each neuron. We then binned each spike train at a temporal resolution of Δ, and consider 1 for active bins and zero for silent bins. Then, we calculated triple-wise interactions (Eq. 4) for each triplet and the average pairwise interaction (Eq. 2) across all pairs within that triplet. To determine the statistical significance of calculated interactions, we compared them with a null distribution of independent spike trains, generated via shuffling ^101,102^. Notably, we derived analytical relations for mean and variance of the pairwise and triple-wise distributions under shuffling (details in the next section). We then identified significant interactions by comparing the observed triplet interactions (⟨ θ_12_ ⟩, θ_123_) with the mean and variance of the shuffled distributions and kept interactions with significant deviations from the shuffled distributions using a high z-score threshold (*z* > 4, p-value < 0.01). A significant interaction parameter thus indicates that the observed joint activity departs from what would be expected under independent spiking activities, beyond finite sampling effects. Our method directly quantifies the structured departure from the null distribution, rather than testing for higher-order interactions relative to a fitted pairwise model ^18^.

### The null distribution for concurrent spike trains, an analytical analysis

We have used the existing correlation in spike trains to calculate pairwise (θ_12_) and triple-wise interactions (θ_123_). The fundamental assumption behind this modelling is that the two uncorrelated neurons generate spike trains that are not correlated at any level, producing zero values for θ_12_ and θ_123_. Therefore, non-zero θ_12_ and θ_123_ are signatures of some level of correlations in the activity of two neurons. To further illustrate this assumption, we produce a simple example of two neurons, each with its independent spiking pattern that obeys Bernoulli distribution: for the *i*-th neuron, the probability of having a spike in each bin is *p*_*i*_ and the probability of being silence is 1 − *p*_*i*_. We can therefore calculate the probability of (1, 1) pattern as *P*_11_ = *p*_1_ *p*_2_; similarly the three other probabilities are calculated as *P*_10_ = *p*_1_(1 − *p*_2_), *P*_01_ = (1 − *p*_1_)*p*_2_, and *P*_00_ = (1 −*p*_1_)(1 −*p*_2_). Using these probabilities, we obtain θ_12_ = log (*P*_11_*P*_00_/(*P*_10_*P*_01_)) = 0. We can easily extend this line of reasoning to three or more neurons. However, there is a practical limitation to obtain the underlying probability, namely, we need to perform a large number of samplings (mathematically speaking, we need an infinite number of samplings). In reality, however, we deal with spike trains of finite length, which compromises the fundamental assumption of independence.

Given two independent spike trains observed over the same duration *t*_obs_, we divided the observation period into *N*_t_ bins of equal width Δ = *t*_obs_/*N*_t_. We have attributed 1 to a bin if one or more spikes have occurred during its time window, and 0 otherwise. Consequently, we obtain two binary sequences of length *N*_t_, consisting of 0 and 1. We know that *n*_1_ (or *n*_2_) occurrences of 1 exist in the first (or second) sequence. This yields *p*_1_ = *n*_1_/*N*_t_ as the probability of spiking in the first spike train, and *p*_2_ = *n*_2_/*N*_t_ for the second one. Following the reasoning in the last paragraph, we try to calculate the probability of finding the (1, 1) pattern as *P*_11_ = *p*_1_ *p*_2_. However, this implies that *p*_1_ *p*_2_*N*_t_ numbers of the (1, 1) pattern should occur in two concurrent sequences. It is easy to see how this is a wrong conjecture: consider two concurrent binary sequences of length *N*_t_ = 10, each with *n*_1_ = *n*_2_ = 5 occurrence of ones, i.e. *p*_1_ = *p*_2_ = 0.5. The aforementioned reasoning means that, regardless of the specific arrangement of the “1”s and “0”s in two sequences, a total of *p*_1_ *p*_2_*N*_t_ = 2.5 patterns of (1, 1) should always emerge; this is evidently wrong. The number of emerged (1, 1) patterns in each instant of observation depends on the detailed permutation of “1”s and “0”s in that case. As we shall see, when averaged over all possible permutations, the average number of emerged (1, 1) patterns is indeed *p*_1_ *p*_2_*N*_t_; but the number of (1, 1) patterns in each observation varies according to the detailed permutation of ones and zeros there. However, these variation means that we cannot always attribute the observed non-zero value of θ_12_ (or θ_123_ for three neurons) to the genuine level of correlation between neurons.

We therefore need a mechanism to determine whether the observed values of θ_12_ and θ_123_ result from a genuine correlation between spiking neurons ^18^, or are simply the side effect of a finite number of samplings. Here, the concept of null distribution is helpful ^103^, which is the probability distribution of θ_12_ (or θ_123_) expected for two (or three) uncorrelated spike trains. By comparing empirical values against this null distribution, we can assess whether observed θ_12_ (or θ_123_) values belong to that distribution, i.e., they are the result of a finite number of sampling, or whether they reflect significant interactions between neurons.

As mentioned above, the most common approach to address this problem is shuffling. We shuffle the binned spike trains 𝒩_sh_ times and compute θ_12_ and θ_123_ after each shuffle. We then calculate the mean and standard deviation of θ_12_ and θ_123_ over 𝒩 _sh_ shuffles and use them to estimate the probability that the empirical values belong to the null distribution. For example, if the difference between empirical value and shuffle mean exceeds four standard deviations (i.e., z-score> 4), the probability that the empirical value belongs to the null distribution is less than 6.3 × 10^−5^. However, the problem with shuffling is its computational cost. Fig. 1d compares the result of 𝒩_sh_ = 100 and 10^6^ shuffles which we have used as ground truth. There is a clear difference between the two, indicating that an adequate number of shuffling is required to obtain reliable estimates. However, this requires substantial computational time. For long spike trains, even 100 shuffles can be prohibitively expensive, especially for neurons in large-scale recordings. For example, the calculation time for 100 permutations of three spike trains with *N*_*t*_= 90060 bins, *n*_1_ = 22500, *n*_2_ = 18000 and *n*_3_ = 13500 is *t*_sh_ = 79 s; this means *t*_sh_ ∼ 2 hour for 10^4^ shuffle, and *t*_sh_ ∼ 9 days for 10^6^ shuffle, whereas the calculations are almost instantaneous with our analytical method.

To circumvent this issue, we suggest a novel approach to analytically calculate the null distribution. Our analytical approach yields a closed form relation for the null distribution, hence providing exact closed form relations for mean and standard deviation of the higher-order interactions. For the ubiquitous limit of long enough spike trains, *N*_*t*_ *≥* 1000, we have used the Stirling large number approximation to reduce the closed form relation to a simple analytical expression. Fig.1d shows that our reduced formula produces the same results as the extremely large numbers of shuffles (𝒩_sh_ = 10^6^), while reducing the computational costs to almost zero. The aforementioned time of *t*_sh_ ∼ 9day, now reduces to *t*_anal_ = 0.007sec, which is almost 100 million times faster. We therefore provide a powerful analytical approach to calculate significant HOIs, without the need to obtain computationally costly shuffled distributions for triplets.

### Probability of synchronous activity for two uncorrelated neurons, *P*_II_(*m*)

To approach the null distribution for θ_12_, we begin with two aforementioned independent spike trains, grouped into *N*_t_ bins. We attribute “1” (or “0”) to each bin, depending on the occurrence (or lack) of a spike in that bin; thus we have two binary sequences of length *N*_t_. There is *n*_1_ (or *n*_2_) occurrence of “1”, hence *N*_t_ − *n*_1_ (or *N*_t_ − *n*_2_) occurrence of “0” in the first (or second) binary sequence. Finally, without any loss of generality, we assume that *n*_1_≥ *n*_2_. To construct the null distribution of θ_12_, we consider all possible permutations of “1” and “0” in two sequences. Since there are no additional constraints on the permutations, all arrangements that preserve *N*_t_, *n*_1_, and *n*_2_ are equally probable.

Given these assumptions, we firstly calculate the probability of synchronous activity. It is the probability that two coincident bins both show 1, i.e. the probability for emergence of the (1, 1) pattern. We define *P*(*m*) as the probability of finding *m* patterns of (1, 1), in two concurrent sequences. On the other hand, we know that the total number of (1, 1) and (1, 0) patterns equals the total number of “1”s in the first spike train; thus, with *m* patterns of (1, 1), there will be *n*_1_ − *m* patterns of (1, 0), and similarly *n*_2_ − *m* patterns of (0, 1), and finally *N*_t_ + *m*− (*n*_1_ + *n*_2_) patterns of (0, 0) (see Fig.1c, top). Therefore, *P*(*m*) is not only the probability of emergence of (1, 1) patterns, but the probability of all other patterns, with the respective number of occurrences.

We define

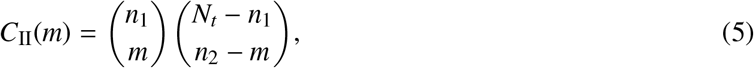

which is the number of possible permutations that produce *m* patterns of (1, 1) and *n*_2_ −*m* patterns of (0, 1). Accordingly, its corresponding probability is the hypergeometric distribution as follows:

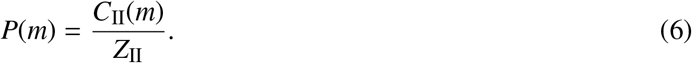

where *Z*_II_ is the normalisation factor, which reads:

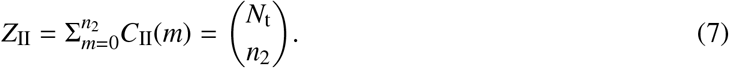

This allows us to calculate the average number of emergence for each pattern, across all permutations:

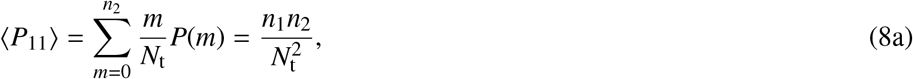

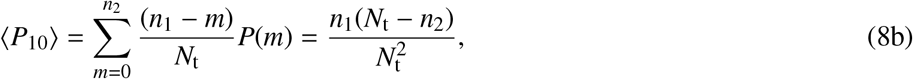

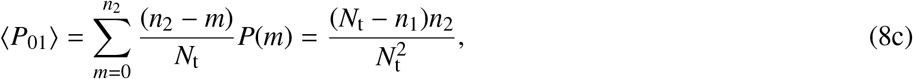

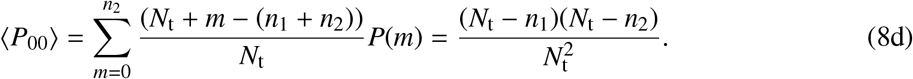

The first and second order moment of θ_12_, using Eq.2 and Eq.6 will be:

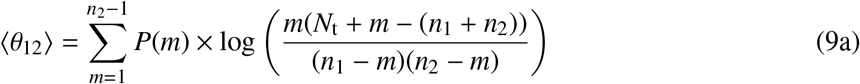

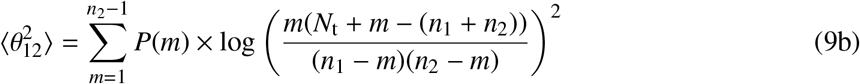

with the standard deviation calculated as 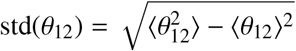.

### The null distribution for θ_12_ in Long-length limit, Stirling approximation

Eq.9a and Eq.9b, provide a closed form relation to obtain ⟨ θ_12_⟩, 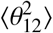, and consequently the variance, var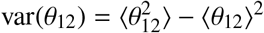, for two independent randomly firing neurons. These closed form relations can safely replace existing shuffling methods which approach the same results through random permutation of binary elements in each sequence, using pseudo-random generators ^104^. As for accuracy, results of Eq.9a and Eq.9b are more accurate, since the shuffling methods always involve some level of error unless we regenerate all possible permutations by running over large number of permutations. However, reliable results can be achieved with a reasonable number of permutations; this is the key advantage of Monte Carlo methods, which come at some computational cost ^105^.

Supplementary figure 1 compares the computational time required to obtain 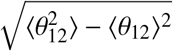, using 10^2^, and 10^6^ number of permutations, with the exact result of Eq.9b. It is evident that the analytic results are much faster, but it is also clear that the timing increases with the sample size (number of total bins in the spike train). So, it is tempting to find a still faster approach for the long spike trains. To this end, we use the Stirling’s approximation for factorial in large numbers ^106,107^:

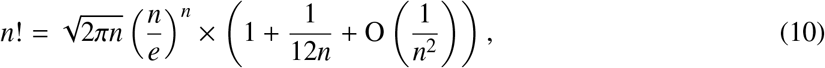

or equally:

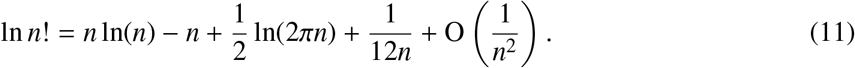

Using Eq.11, and after a detailed calculation, we simplify Eq.6 to its asymptotic limit of:

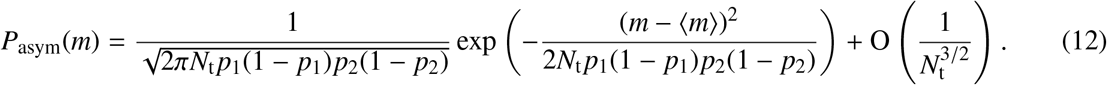

for large *N*_t_, where *p*_1_ = *n*_1_/*N*_t_ and *p*_2_ = *n*_2_/*N*_t_. Figure 1d, compares this asymptotic relation with the exact result of Eq.6.

Using *P*_asym_(*m*), we calculate ⟨ θ_12_ ⟩, 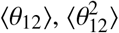. To this end, we use the method of Saddle point approximation for Gaussian expectation values. It allows us to calculate the average of any function *f* (*m*), where *m* obeys a Gaussian distribution, as:

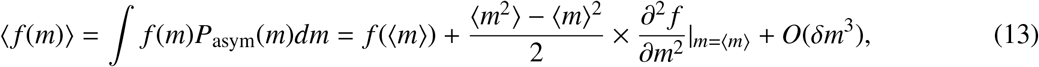

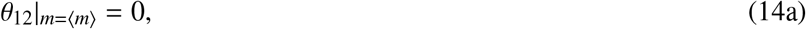

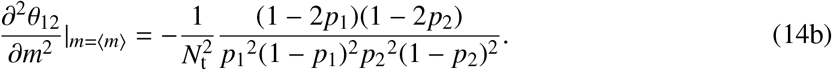

We therefore obtain:

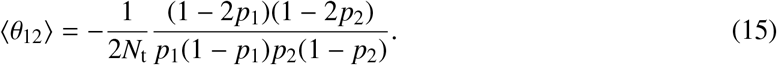

Also,

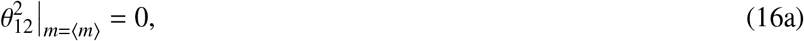

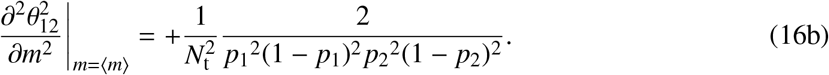

which results in:

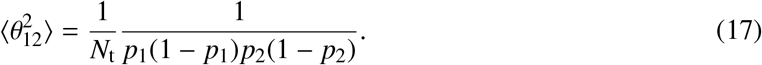

In many practical cases, Eq.15 and Eq.17 replace a time-consuming procedure with an efficient and reliable answer. For example, for two spike trains with total length *N*_t_ = 10 000 bins, one has *n*_1_ = 2540 and the other *n*_2_ = 1450 spikes, the exact answers read ⟨θ_12_⟩ = 7.45 × 10^−4^ and 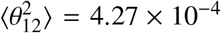, whereas the asymptotic ones are ⟨θ_12_⟩ = 7.44 × 10^−4^ and 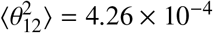, which shows an error less than 1%.

### Probability of activity patterns for three uncorrelated neurons, *P*_III_

As previously discussed, when *N*_t_, *n*_1_, and *n*_2_ are known and fixed, knowing the number of (1, 1) pattern (i.e. *m*) allows us to determine the number of three other patterns, (1, 0), (0, 1), and (0, 0). The reason is that fixed values of *N*_t_, *n*_1_, and *n*_2_ produce three independent conditions; knowing *m*, we will have the fourth condition, allowing us to calculate four unknown numbers of patterns. Firstly, the total number of four patterns is *N*_t_, i.e., the total number of bins. Moreover, the number of (1, 1) and (1, 0) patterns together is the total number of all bins in which the first neuron generates a spike, i.e. *n*_1_. Similarly, the number of (0, 1) and (1, 1) patterns, together, is *n*_2_. Consequently, we can easily calculate *n*_1_ − *m* as the number of (1, 0) patterns, and *n*_2_ − *m* as the number of (0, 1) patterns. Finally, the number of (0, 0) patterns will read as *N*_t_ + *m* − (*n*_1_ + *n*_2_).

For three neurons, however, we know four initial values of *N*_t_, *n*_1_, *n*_2_, and *n*_3_; on the other hand, the three neurons produce 2^3^ possible patterns (see Fig.1c, bottom). Therefore, we need four other independent variables to be able to calculate the number of times all 8 patterns appear. As for two neurons, we use *m* as the number for all (1, 1) patterns, irrespective of what is happening to the third neuron, i.e. *m* is the number of (1, 1, 1) and (1, 1, 0) patterns altogether. Then, we define *s*_11_ as the occurrence number of the particular (1, 1, 1) pattern. Thus *s*_11_ counts how many times the third neuron generates a spike, while the first two neurons show the (1, 1) pattern. Following this terminology, we define *s*_10_, *s*_01_ and *s*_00_ as the number for 101, 011, and 001 patterns, respectively. However, these would be five variables, four *s* and one *m*; yet, we have an extra condition:

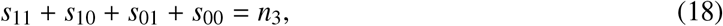

therefore, one variable *s*_00_ = *n*_3_ − *s*_11_ − *s*_10_ − *s*_01_ is not an independent variable. We are left with *s*_11_, *s*_10_, *s*_01_, and *m* as the four independent variables to (I) describe the number of occurrences of each of the four patterns, and (II) calculate their respective probability.

For the case of two neurons, the number of 101 and 100 patterns together, is *n*_1_ − *m*. As *s*_10_ further measures the number of 101 patterns, we will simply have the *n*_1_ − *m* − *s*_01_ number of 100 patterns. A similar reasoning yields *n*_2_ −*m−−s*_01_ as the emergence number of the 010 pattern, and *N*_*t*_ − (*n*_1_ +*n*_2_)+*m* −*s*_00_ as the number of the 000 pattern.

To address the probability, we extend the two neuron’s permutation number, *C*_II_(*m*), with a correcting factor:

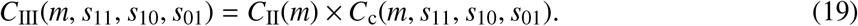

The correcting factor reads:

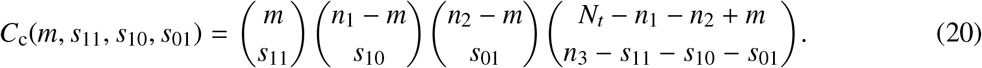

Accordingly, the probability density function for having such patterns (synchronisation probability) reads:

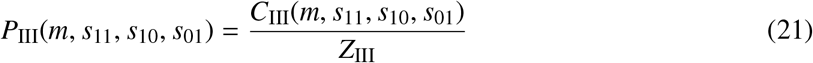

where

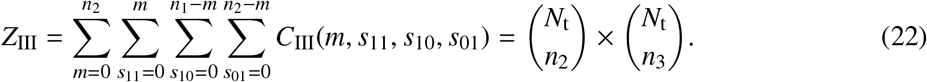

This result comes from a simpler summation of:

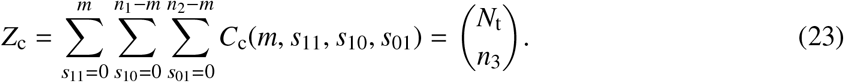

Intuitively, Eq.23 shows that, considering the various permutations of the third binary sequence, we have 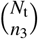 different permutations for each given state of the first and second sequences.

In a similar fashion as in the two neuron’s case, we can calculate the mean probabilities for eight possible patterns 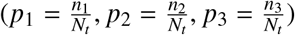:

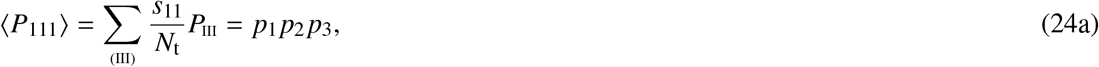

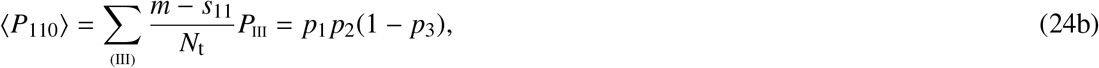

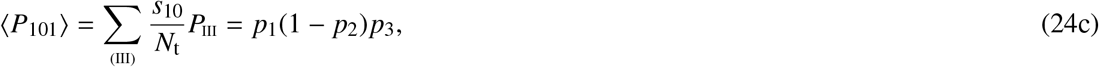

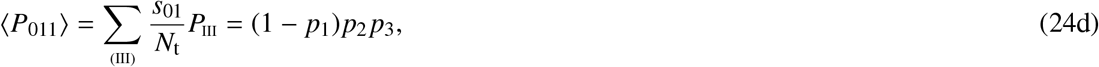

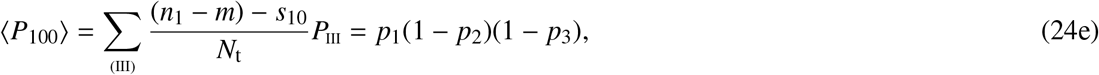

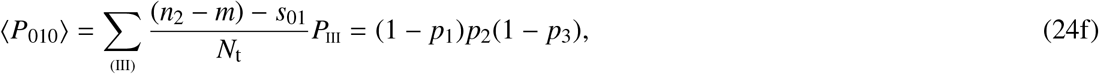

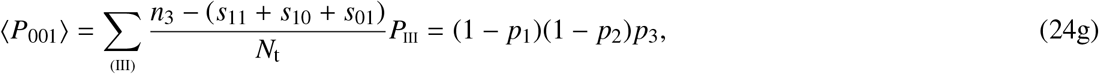

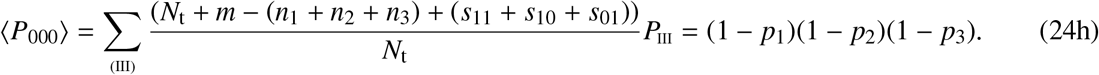

where 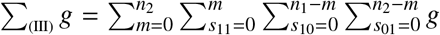 goes over all allowed values of *m, s*_11_, *s*_10_, and *s*_01_, c.f. Eq.22. Similar to the two neurons case, the mean and variance of triple-wise interactions can be calculated as:

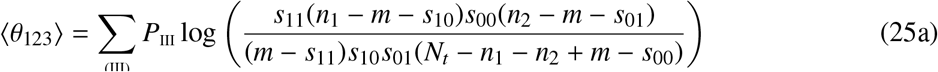

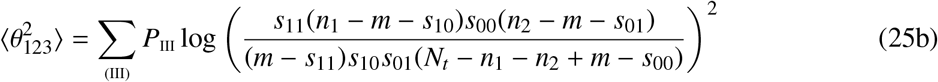

with the standard deviation std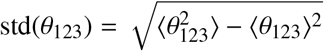.

The analytic relation for the mean and variance of triple-wise interactions of shuffled spike trains helps us to find the exact values rather than simulating a large number of permutations numerically, which will come at a high computational cost. However, similar to the pairwise case, as the total number of bins (length of spike trains) increases, the analytic results become harder to achieve due to the computational cost ((Eq.25a and Eq.25b)). For such a case, we approximate it for long total bins (*N*_*t*_ > 1000) as an asymptotic relation that is instantaneous.

### Null distribution of θ_123_ in the long length limit

Using the saddle point approximation (Eq.13), we can calculate ⟨θ_123_⟩ and 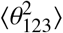. First, using Eq.21, we calculate the saddle point values (⟨*m*⟩, ⟨*s*_11_⟩, ⟨*s*_10_⟩, ⟨*s*_01_⟩):

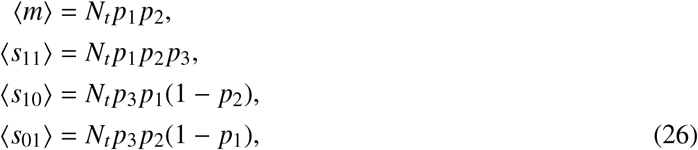

then the covariance matrix of parameters can be calculated as:

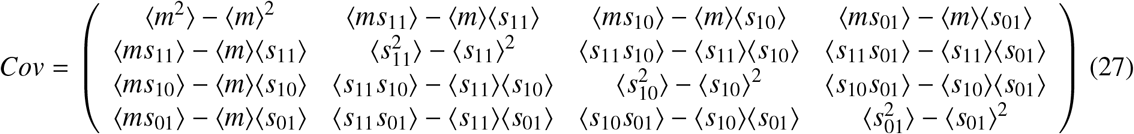

The matrix elements of Eq.27 can be obtained using Eq.21:

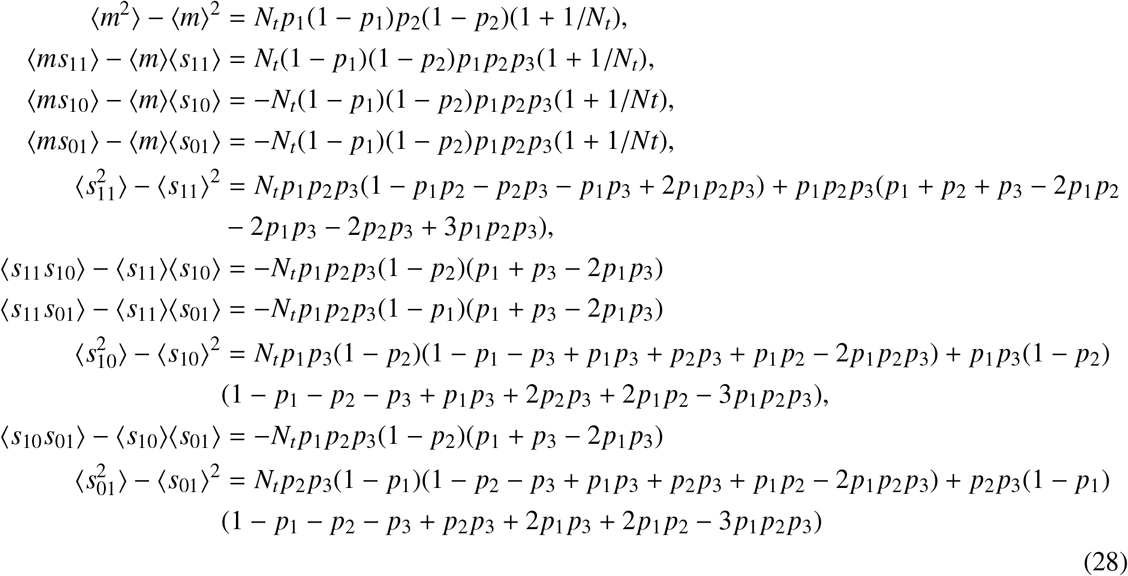

We can calculate the average triple-wise interactions and its corresponding variance around the saddle point (*m* = ⟨*m*⟩, *s*_11_ = ⟨*s*_11_⟩, *s*_10_ = ⟨*s*_10_⟩, *s*_01_ = ⟨*s*_01_⟩). The final result reads:

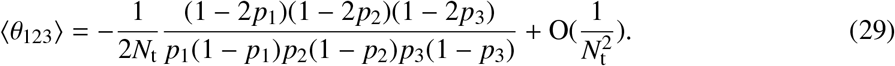

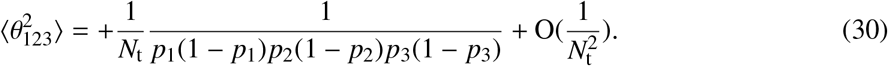

where the standard deviation is std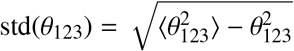. Figure 1d (bottom) compares the results of our asymptotic relations (Eq.29 and Eq.30, green curves), with that of one million rounds of shuffling (red points/curve), and with results of 100 shuffles (sky blue points/curve), and with the closed form analytic series in Eq.25 (black curve). For total bins of less than 1000, the analytic series perfectly match with 10^6^ shuffles, but we cannot go higher than *N*_t_ = 1000 due to its high computational cost. Thus, for longer spike trains, we use the asymptotic relations of Eq.29 and Eq.30. There is a good agreement between the asymptotic results and the results obtained from one million shufflings, and this agreement continuously improves as *N*_t_ increases. Shuffling for 100 times takes much less computational time (compared to 10^6^ shuffling), but it shows a non-negligible error evident in the deviation from all other curves; this deviation does not necessarily decrease with increasing *N*_t_. The asymptotic relation is far faster than other approaches (almost instantaneous), produces a negligible error and its error further reduces as *N*_t_ increases. To get a sense of its error, for the specific number of bins *N*_t_ = 10000, we have investigated a wide range of firing rates of 0.1 ≤*p*_*i*_ *≤*0.9, with a step size of 0.1. We have compared the result of the asymptotic relation with that of 10^6^ permutations for these 9^3^ = 729 points, and computed the relative error. The relative error reaches as high as 3% for a few points, but remains less than 1% for the majority of them. As the relative error should behave like 1/*N*_t_, we further conclude that for *N*_t_ > 30000, the relative error will be less than 1% for all the probed points.

### Guide map of neural interactions

We previously linked neuronal pairwise and higher-order interactions to basic motifs in the interaction plane ^50^ and provided a guide map of neural interactions. This map consists of regions, each associated with a specific circuit motif ^50^(Fig. 3a). Each region corresponds to a distinct motif of shared inputs between three neurons. For example, the grey region represents excitatory common input to all three neurons. Each region is delineated by two boundary lines. The first boundary line corresponds to the regime of strong common input amplitudes given to postsynaptic neurons. The second boundary line is derived by fixing the spontaneous activity of postsynaptic neurons at a low rate (i.e. 1Hz) while systematically increasing the common input amplitude from zero until it saturates at the strong amplitude boundary ^50^. Therefore, the region contains the results of pairwise and triple-wise interactions for a range of spontaneous activity (1 −100 Hz) and common input’s amplitude (from 0 to strong amplitudes). The merit of this guide map in the plane of triple-wise versus pairwise interactions is that it can distinguish among circuits that cannot be differentiated by pairwise interactions alone. By superimposing the interactions calculated from empirical data onto this theoretical guide map, the underlying (or the reduced) shared circuit motifs are revealed and can be quantitatively characterised.

### Defining behavioural states

We applied the speed threshold of 2 cm/s to separate trials and label them as stationary (speed < 2 cm/s) and running trials (speed > 2 cm/s). We selected mice that maintained the speed threshold for at least 4 continuous stationary and running trials to ensure reliable estimation of interactions due to change in the state. For each neuron, we concatenated continuous trials of the same type (stationary or running) together. To ensure a comparable analysis, we aimed to include roughly equal numbers of trials for stationary and running states, thus maintaining similar spike train lengths for both groups. We included only animals that produced more than 30 significant interaction points in both stationary and running trials. This approach allowed us to examine how speed influences neural interactions across different brain regions and brain states.

### Simulation of balanced networks of excitatory and inhibitory integrate-and-fire neurons

We simulated randomly connected networks of leaky integrate-and-fire (LIF) neurons, comprising 800 excitatory and 200 inhibitory neurons ^64^. The membrane potential dynamics of each neuron are governed by:

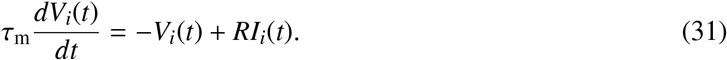

where *I*_*i*_(*t*) is the total synaptic current of all spikes that arrive at postsynaptic neuron *i* at time *t*. Spikes are modelled as delta functions in LIF neuron model:

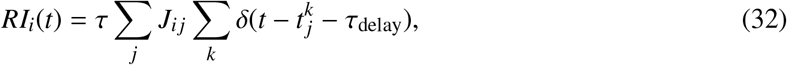

where *J*_*i−j*_ is the synaptic weight from neuron *j* to neuron *i* and 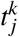 denotes the *k*-th spike time of presynaptic neuron *j*. The membrane time constant was τ_*m*_ = 20 ms, the refractory period was τ_*re−f*_ = 2 ms, and synaptic delay time was τ_delay_ = 1.5 ms. The resting voltage was set at *V*_rest_ = 0 mV. When the membrane potential reached the threshold *V*_θ_ = 20 mV, the neuron generated a spike and reset to *V*_reset_ = 10 mV. Neurons were sparsely connected with connection probability *c*_*p*_ = 0.1, yielding approximately 100 inputs per neuron. Excitatory and inhibitory synaptic weights were set to *w*_*e*_ = 0.1 and *w*_*i*_ = −*gw*_*e*_, respectively, where g is the ratio of inhibitory to excitatory synaptic strengths and typically ranges from 4 to 8 in cortical networks, with *g* = 4 − 5 marking the transition to asynchronous irregular activity in the these networks ^64^. Each neuron received an external Poisson input with a rate of ν_ext_ = 2.5 Hz to drive spontaneous activity. The network was simulated using the Euler method with time step δ*t* = 0.1 ms for *T* = 100 s. We set *g* = 5 to investigate the asynchronous irregular regime of cortical activity. We also varied the ratio g and external drive rate ν_*ext*_ to explore different dynamical regimes, from asynchronous irregular regimes of spiking activity to synchronous oscillatory states ^64^, allowing us to examine how interaction motifs emerge across distinct network activity patterns.

### Ethics statement

This study is entirely computational and involves no human participants, animal subjects, or personal data. No ethical approval was required.

## Data availability

All data used in this study are publicly available online. Sources and links are provided in the references.

## Acknowledgments

This work is supported by the Wellcome Trust (award ref. 225412/Z/22/Z to S.S.). We thank Elizabet de Laittre, Harrison Maddox, and Marco Ferreti for critical reading of the manuscript and all members of the Sadeh lab for their feedback. H.S. is supported by JSPS KAKENHI Grant Number JP 24K21518, 25K03085.

## Competing interests

The authors declare no competing interests.

## Notes

### Competing Interest Statement

The authors have declared no competing interest.

