## Supplementary Information for "Inferring state-dependent functional circuit motifs using higher-order interactions analysis"

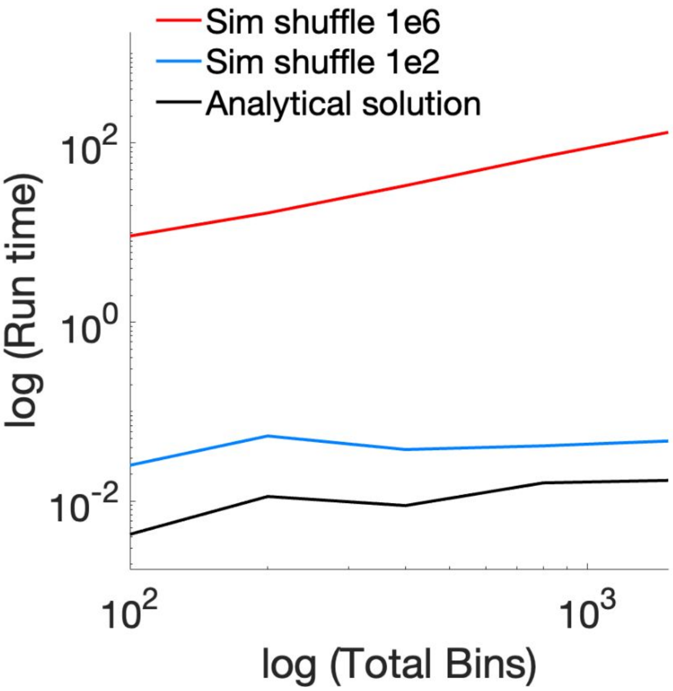


**Supplementary Fig 1.** Comparing the total time for calculating the mean and variance of pairwise interactions using shuffling by simulation (100 times and $10^6$ times) with analytic calculation (Methods). The graph shows the time (s) versus total bins of spike trains. The firing of two neurons are 25 and 20 Hz.

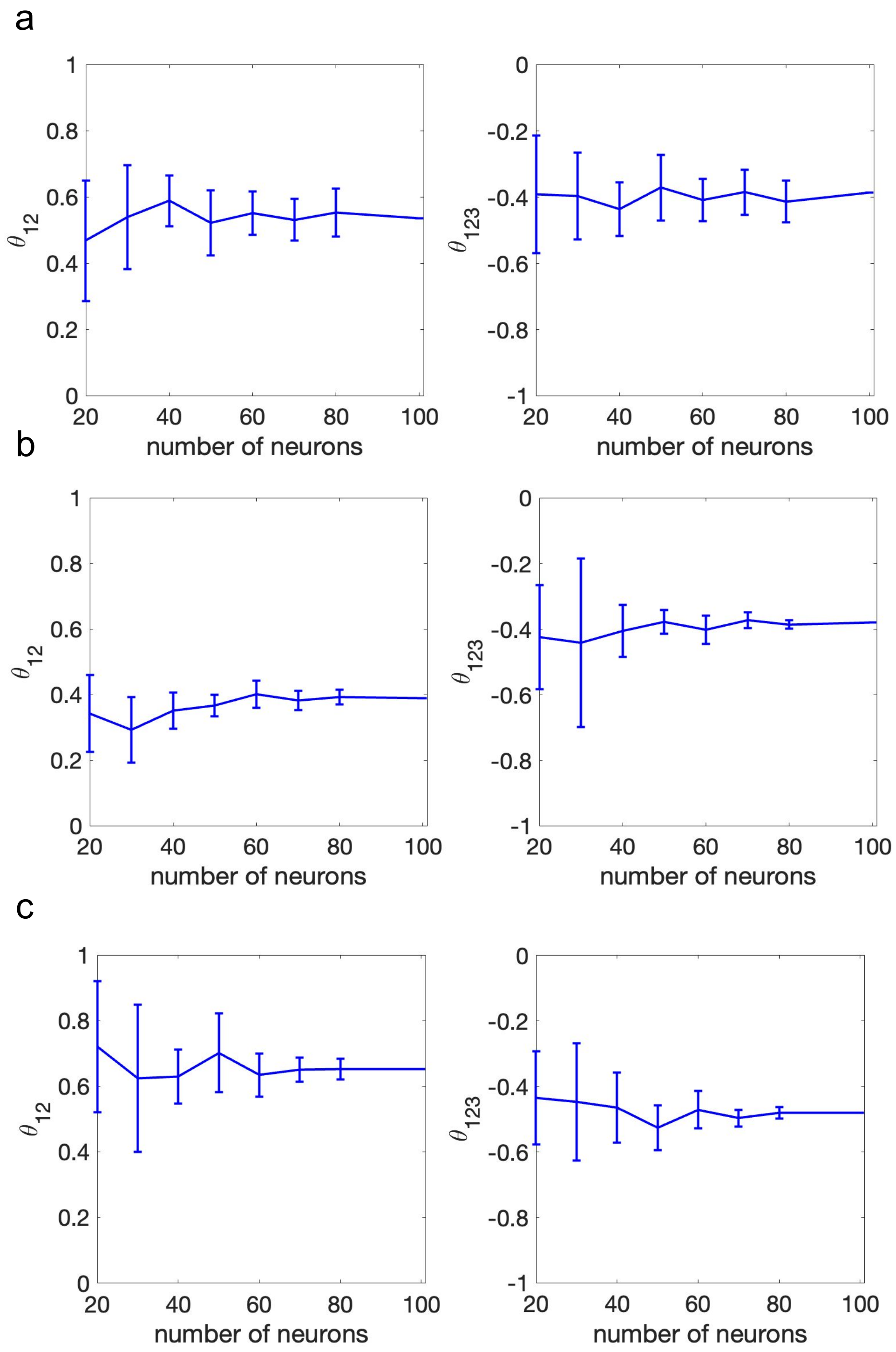


**Supplementary Fig 2.** Dependence of neural interactions on population size for three mice (a, b, and c) in Functional connectivity dataset. The pairwise (left) and triple-wise (right) interactions were quantified for randomly sampled subsets of neurons from a population of 100 units. Data points represent mean interaction strengths, with error bars indicating standard deviation obtained from 10 independent sampling iterations. The bin size is 20 ms.

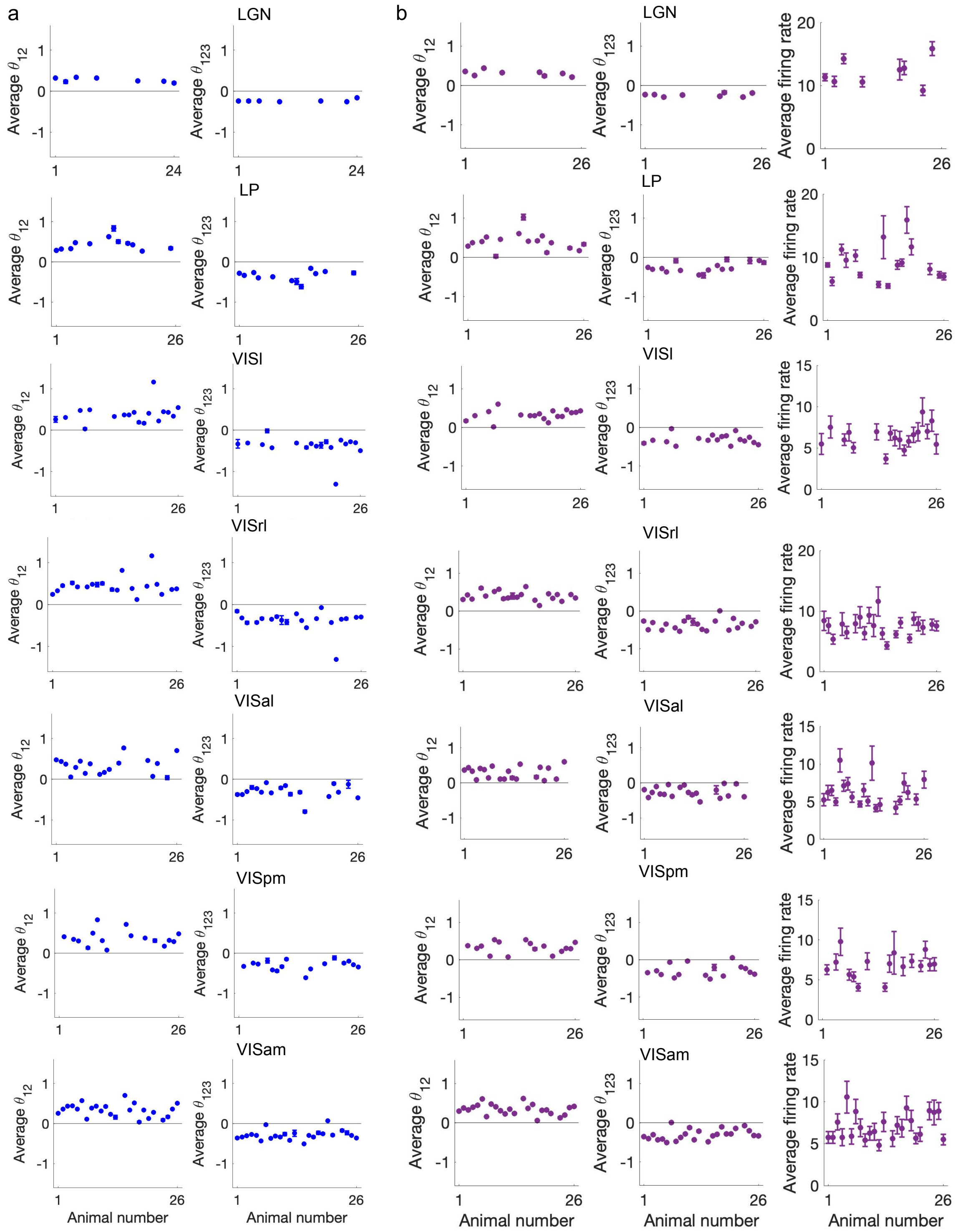


**Supplementary Fig 3.** Average of significant pairwise (left) and triple-wise (middle) interactions for all mice in thalamus nuclei (LGN and LP) and visual hirarchies (VISl, VISrl, VISal, VISpm and VISam) in Functional connectivity dataset for bin size $\Delta$ = 20ms (a), and $\Delta$ = 50ms (b). Firing rates of neurons participating in significant interactions (right). Error bars are SEM, z-score= 4, and p-value< 0.0001. All data shown here are from a functional connectivity dataset.

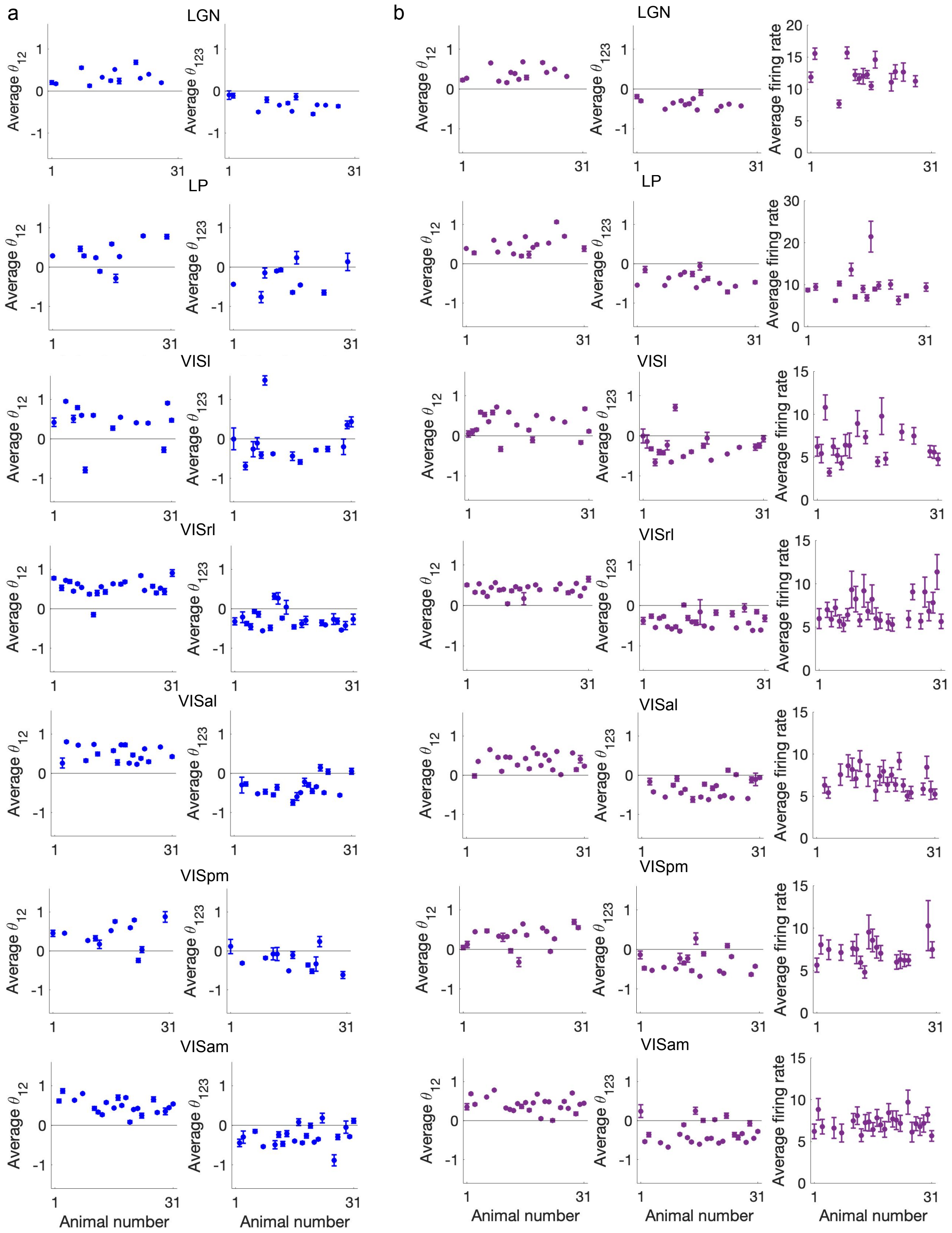


**Supplementary Fig 4.** Average of significant pairwise (left) and triple-wise (middle) interactions for all mice in thalamus nuclei (LGN and LP) and visual hirarchies (VISl, VISrl, VISal, VISpm and VISam) in brain observatory dataset for bin size $\Delta$ = 20ms (a), and $\Delta$ = 50ms (b). Firing rates of neurons participating in significant interactions (right). Error bars are SEM, z-score= 4, and p-value< 0.0001. All data shown here are from a Brain Observatory dataset.

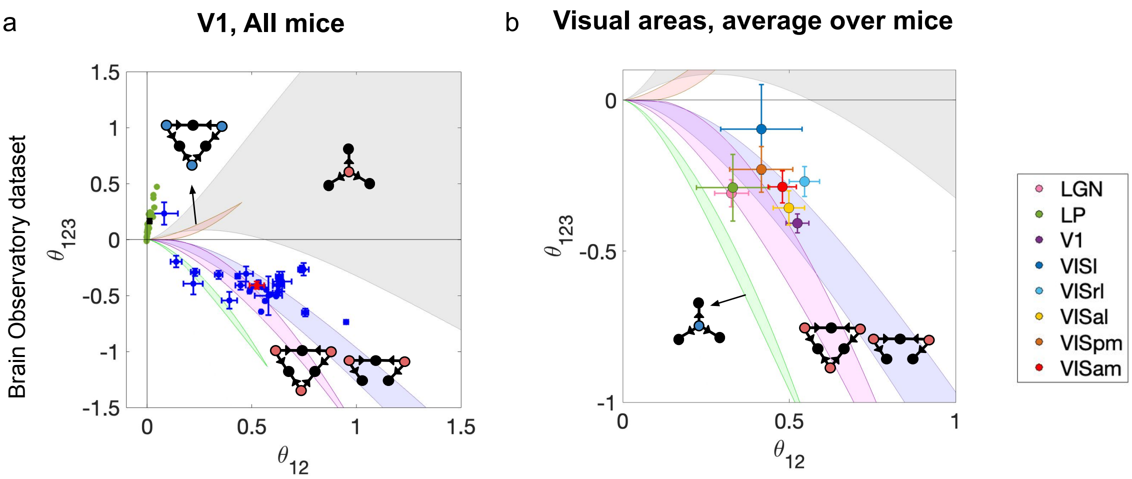


**Supplementary Fig 5. a** The interactions of each individual mouse V1 data (blue) and the average over all mice (red) on the guide map of neural interactions (Bin = 20 ms). The green dots are interactions from shuffled spike trains and the black dot shows the mean±SEM. **b** Average pairwise and triple-wise interactions across thalamus and visual areas (from LGN to VISam, Brain Observatory datasets) plotted on the guide map. Error bars represent SEM.

a

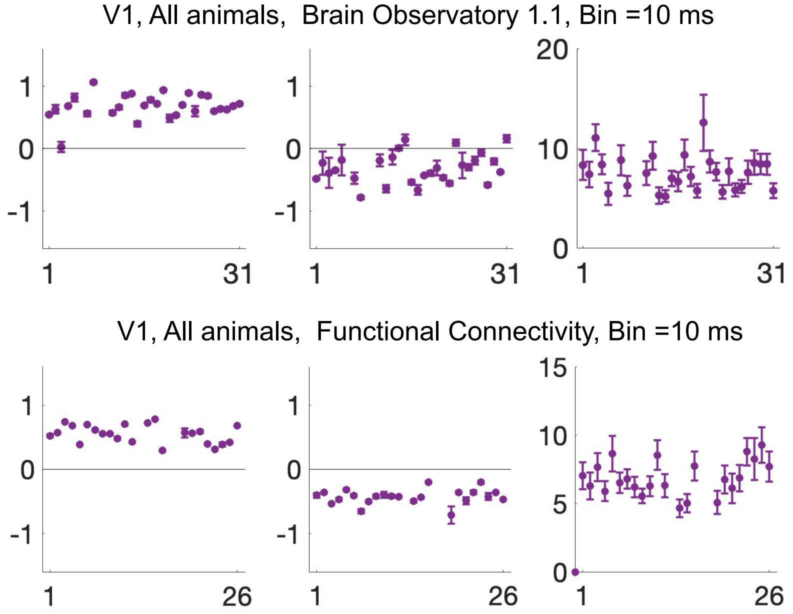


b

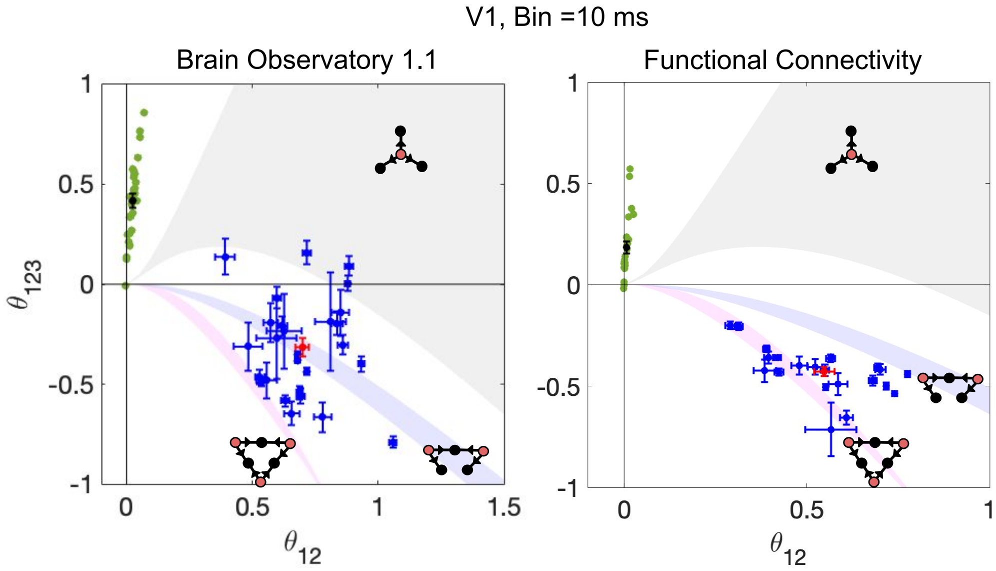


**Supplementary Fig 6. a** Average of significant pairwise (left) and triple-wise (middle) interactions of V1 versus animal's number for each mouse, when bin size is 10 ms (p-value < 0.0001). Firing rates of neurons participating in significant interactions (right). **b** The average pairwise and triple-wise interactions for each mouse (blue dots) and the average across all mice (red) are shown for each region on the guide map of neural interactions. The Bin size is 10 ms and error bars are SEM. The green dots are interactions from shuffled spike trains and the black dot shows the mean ± SEM.

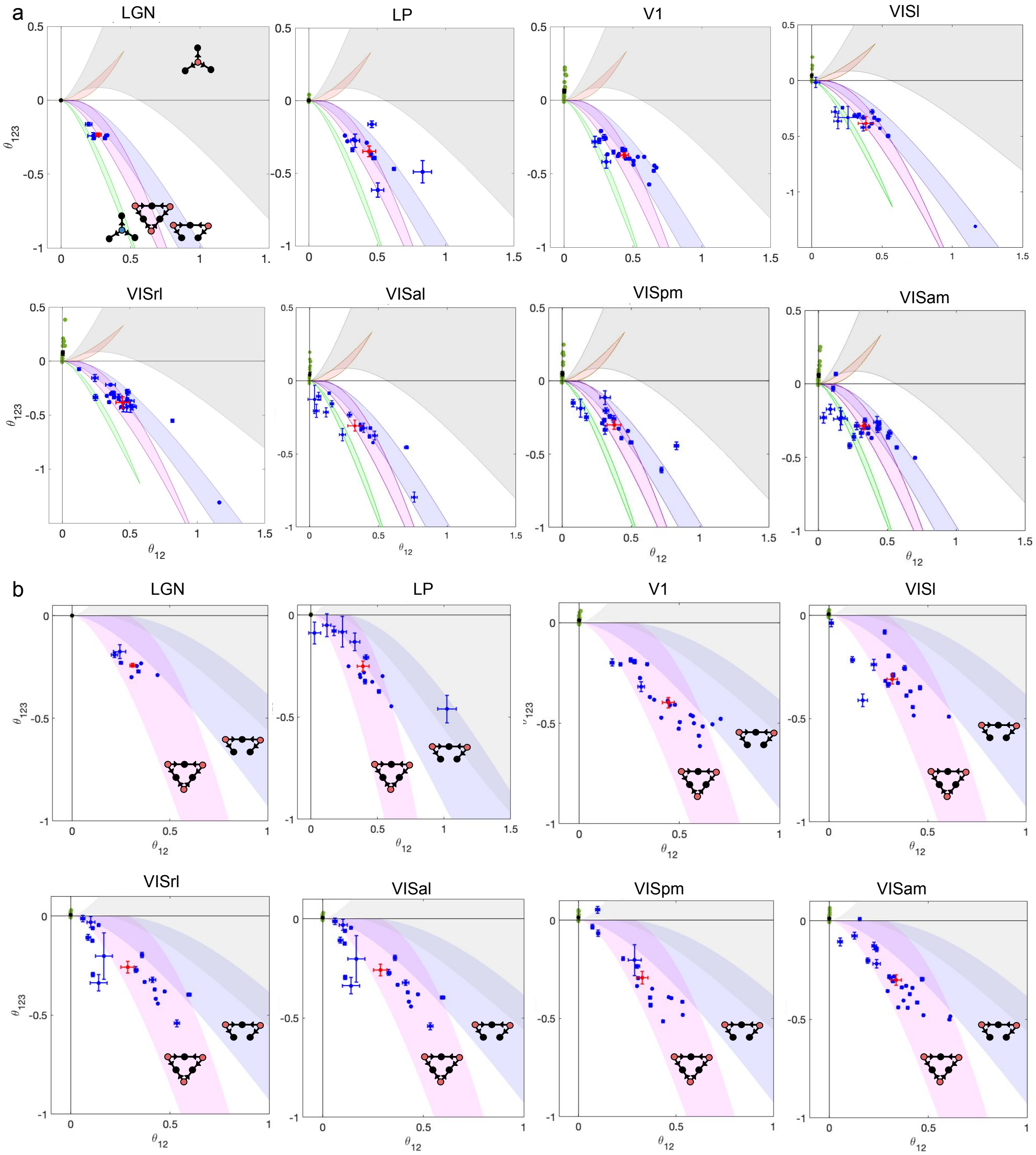


**Supplementary Fig 7.** The average pairwise and triple-wise interactions for each mouse (blue dot) and the average of all mice (red) are shown for each region across visual hierarchy on the guide map of neural interactions. The pink region is associated with symmetric excitatory-to-pairs, violet region is for asymmetric excitatory-to-pairs and the gray region is associated with excitatory-to-trio motifs. Regions are LGN, LP, V1, VISl, VISrl, VISal, VISpm and VISam (Functional Connectivity dataset). The time window is $\Delta = 20\,ms$ in a and $\Delta = 50\,ms$ in b. The error bars are SEM. The green dots are interactions from shuffled spike trains and the black dot shows the mean ± SEM.

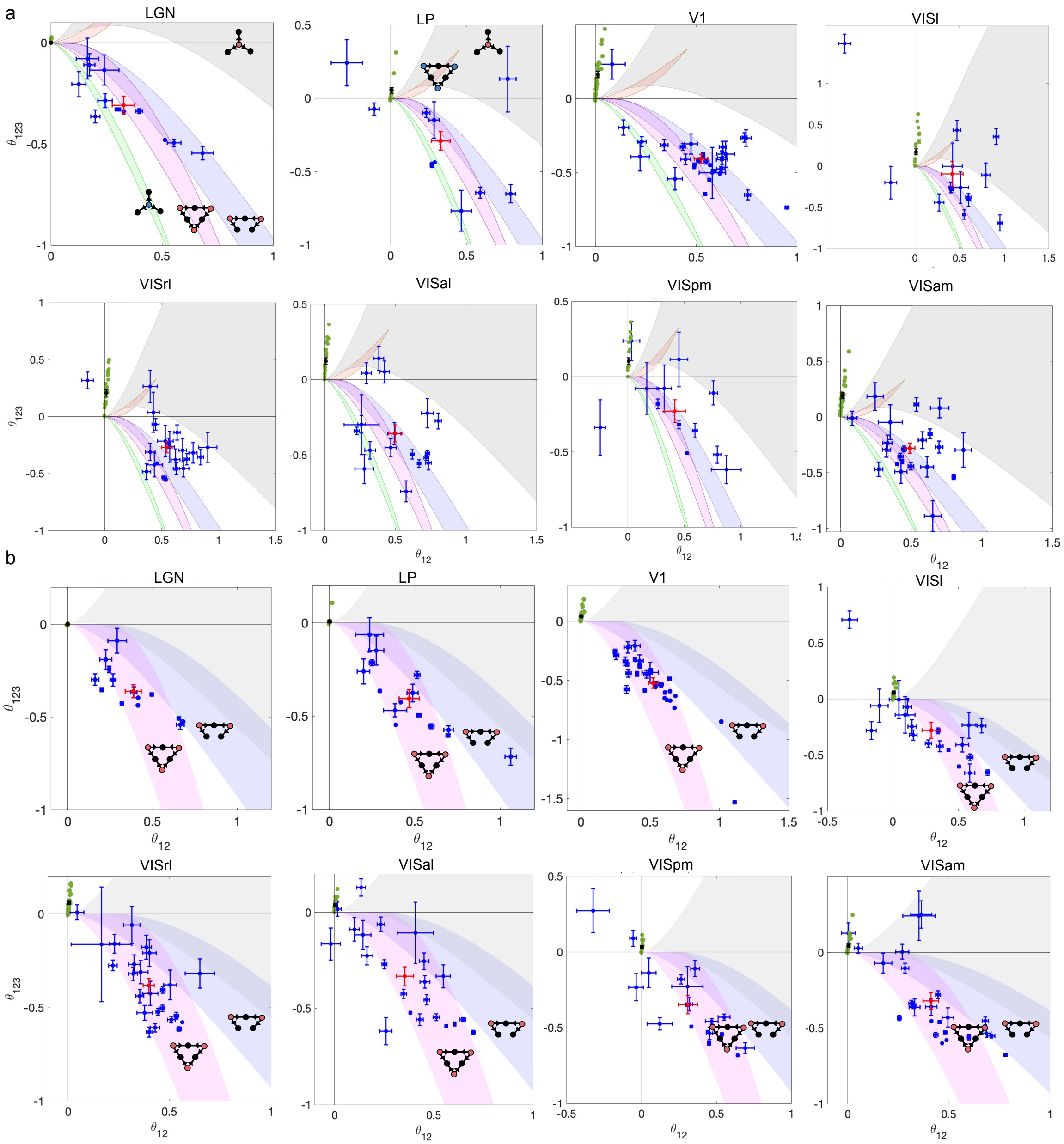


**Supplementary Fig 8.** The average pairwise and triple-wise interactions for each mouse (blue dot) and the average of all mice (red) are shown for each region across visual hierarchy on the guide map of neural interactions. The pink region is associated with symmetric excitatory-to-pairs, violet region is for asymmetric excitatory-to-pairs and the gray region is associated with excitatory-to-trio motifs. The green region is associate with inhibitory to trio motif and the orange region is for inhibitory to pairs motif. Regions are LGN, LP, V1, VISl, VISrl, VISal, VISpm and VISam (Brain Observatory 1.1 dataset). The time window is $\Delta = 20ms$ in a and $\Delta = 50ms$ in b and error bars are SEM. The green dots are interactions from shuffled spike trains and the black dot shows the mean ± SEM of shuffled spike trains.

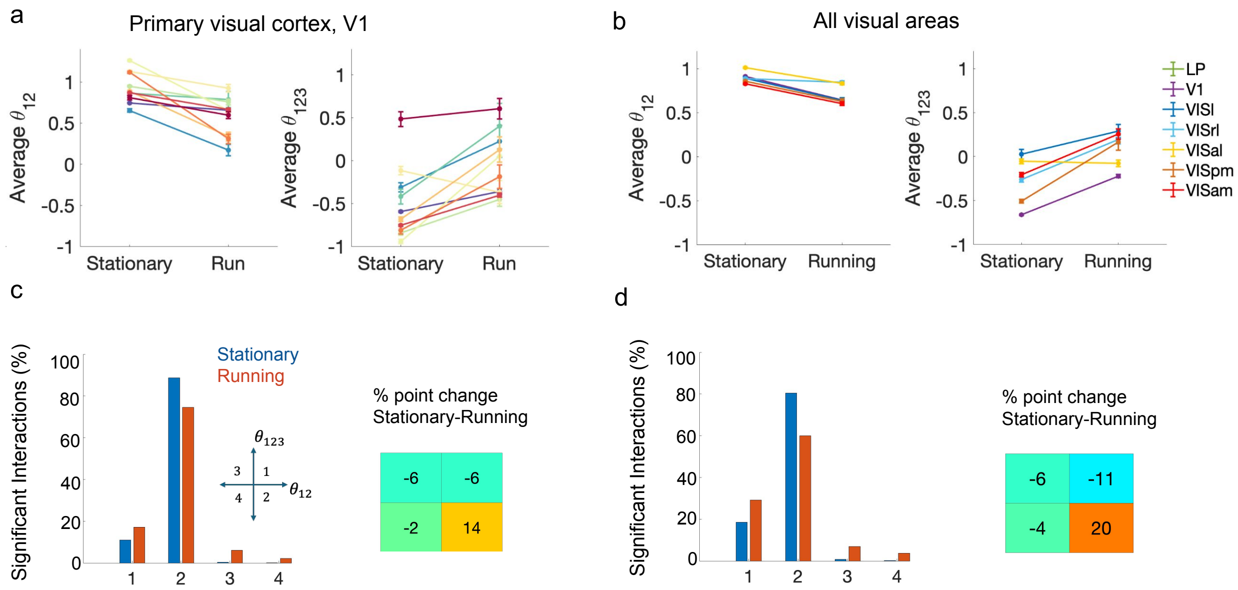


**Supplementary Fig 9.** Higher-order interactions differ in stationary and running states. **a** Average pairwise (left) and triple-wise (right) interactions for stationary and running trials of each mouse in primary visual cortex (V1). Pairwise interactions in nine out of ten animals and triple-wise interactions in eight out of ten animals differ significantly between states. However, firing rates between the two states showed no significant difference in 6 of 10 animals (Suppl. Fig. 10; t-test, $p > 0.05$), with the remaining 4 animals showing significantly elevated rates in [state name] ($p < 0.01$). **b** Pairwise (left) and triple-wise (right) interactions averaged over mice across visual hierarchies. Except for triple-wise interactions in VISal, all pairwise and triple-wise interactions differed significantly between stationary and running states across visual regions. The firing rates show no significant difference for two regions (VISl and VISpm) out of six, between the two states (Suppl. Fig. 10, ttest, $P > 0.05$). Error bars in **a** and **b** represent SEM. The firing rates of neurons participating in significant interactions in **a** and **b** are shown in Suppl. Fig. 10. **c** Left: percentage of significant interactions for stationary and running states in each quadrant of triple-wise versus pairwise plane, in V1. Right: percentage point change between stationary and running data in each quadrant. **d** Left: Percentage of significant interactions for stationary and running states in each quadrant of the interaction plane (triple-wise versus pairwise plane), averaged across all visual areas (left). Right: percentage point change between stationary and running data in each quadrant across all visual areas. The results shown here are from Brain Observatory dataset for bin width 20 ms.

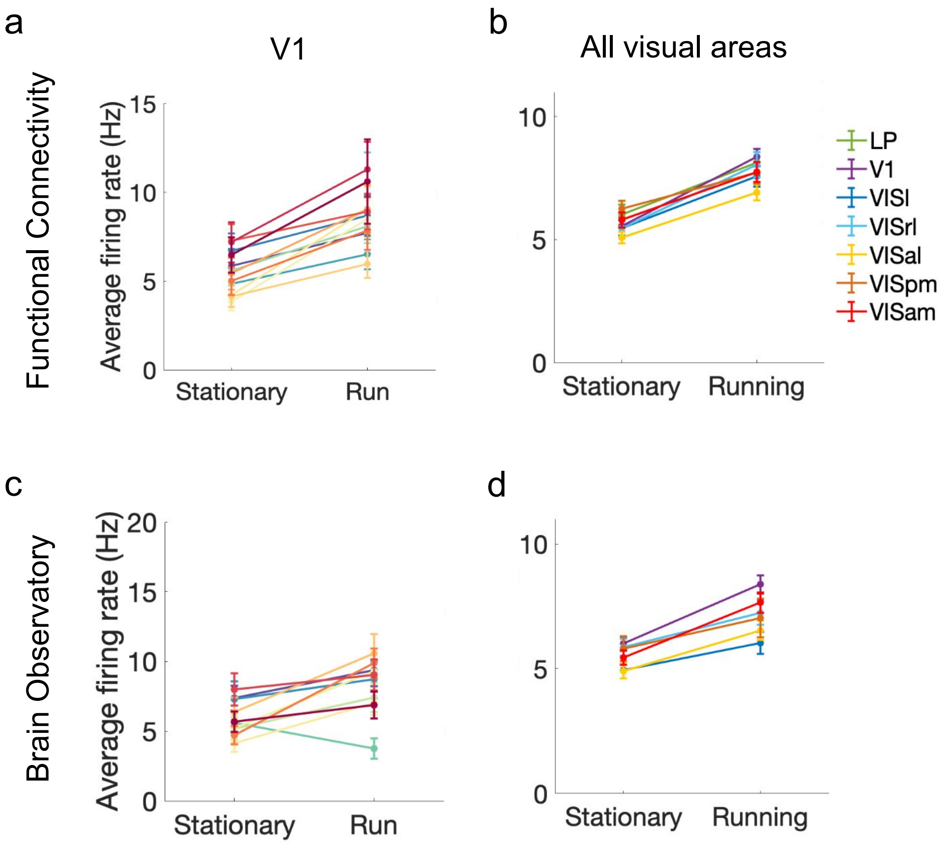


**Supplementary Fig 10.** Firing rates of stationary and running states for each mouse in V1 (left) and across visual areas (right) for two datasets. **a** The firing rates in V1 in six out of thirteen animals show no significant difference between stationary and running states (t-test, $p > 0.05$). **b** All regions show significant difference between the two states. **c** Six out of ten animals show no significant difference of firing between stationary and running states (t-test, $p > 0.05$). **d** Two visual regions VISl and VISpm out of six regions show no significant difference in firing between stationary and running states. The bin size in all is 20 ms.

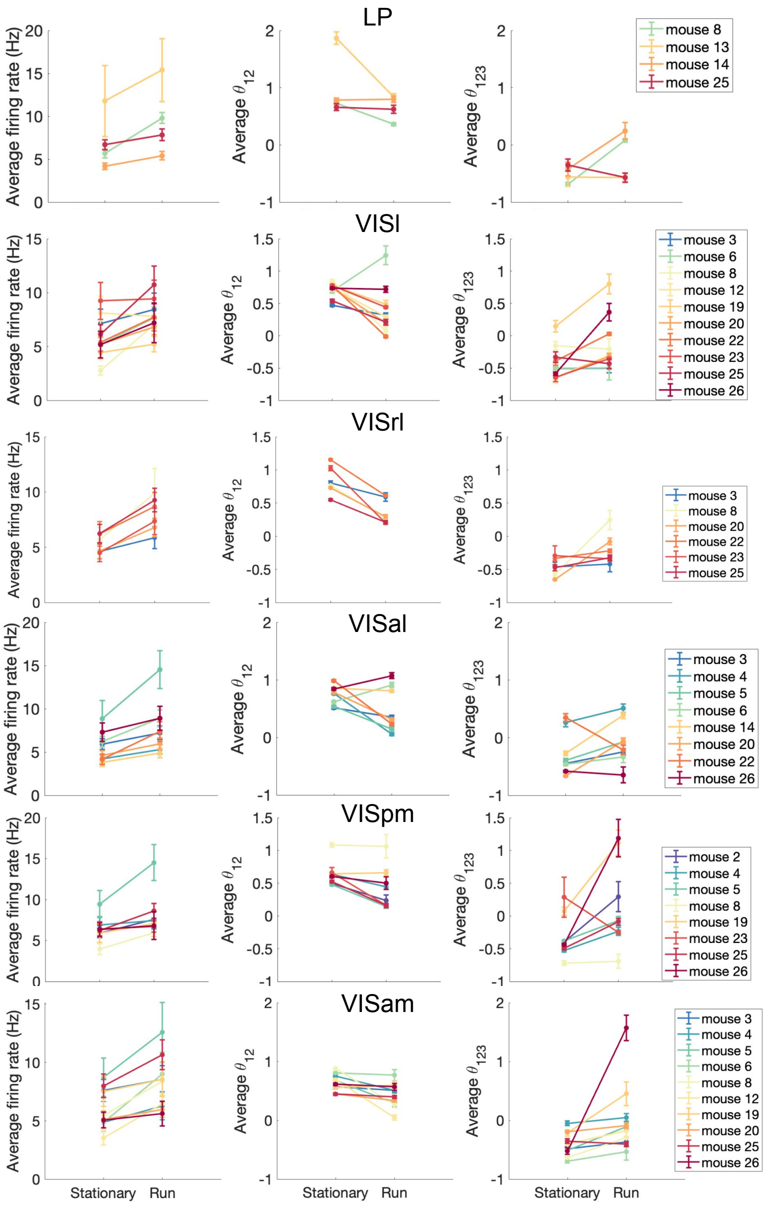


**Supplementary Fig 11.** Average firing rates, pairwise and triple-wise interactions between stationary and running states across visual hierarchies, from LP to VISam. In LP, firing rates, pairwise, and triple-wise interactions in two out of four mice show no significant difference between states (t-test, $p > 0.05$). In VISl, Firing rates in seven out of ten animals doesn't show significant interactions, while pairwise interactions in one animal and triple-wise interactions in four animals show no significant difference between stationary and running. In VISrl, firing rates in four out of six animals show no significant difference between states, while pairwise interactions in zero animals and triple-wise interactions in two animals show no significant difference. In VISal, firing rates in six out of eight animals didn't show significant interactions. However pairwise in one and triple-wise in two animals show no significant interactions. In VISpm, firing rates in seven out of eight animals show no significant difference; however, pairwise in two animals and triple-wise in one animal show no significant difference. In VISam, firing in eight out of ten animals, while pairwise and triple-wise in five animals show no significant difference between stationary and running states (t-test, $p > 0.05$). Data shown here are from Functional connectivity dataset and the time window is $\Delta = 20$ms.

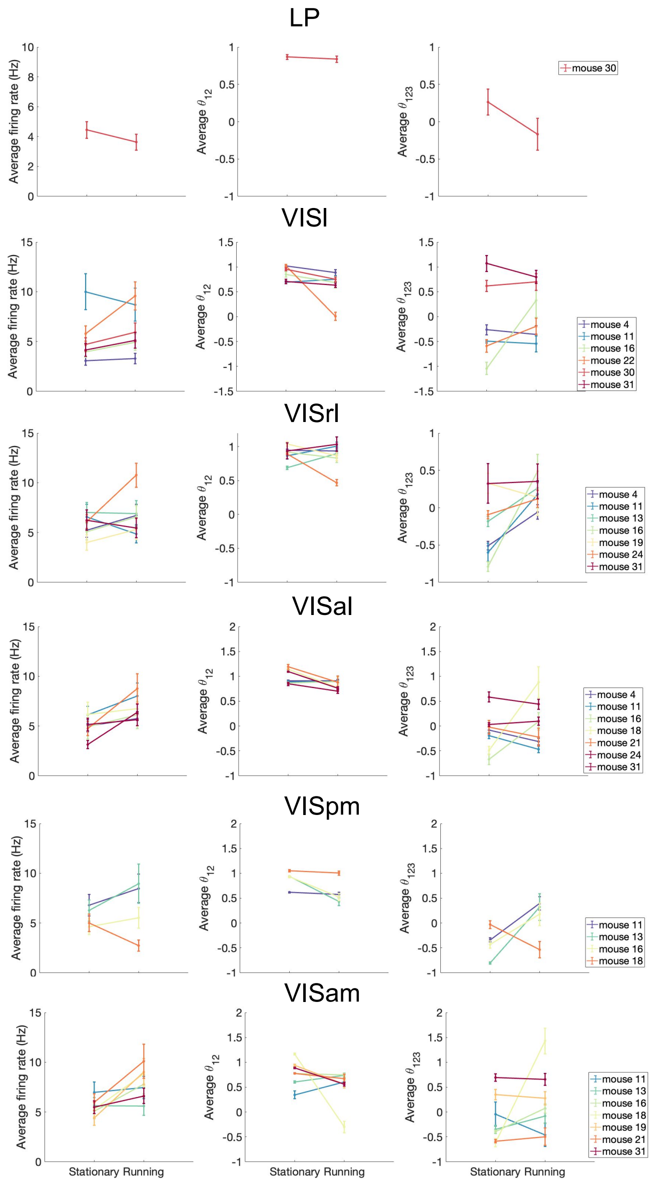

**Supplementary Fig 12.** Average firing rates, pairwise and triple-wise interactions between stationary and running states across visual hierarchies, from LP to VISam. In LP, firing rates, pairwise, and triple-wise interactions show no significant difference between states in the animal (t-test, $p > 0.05$). In VISl, Firing rates in five out of six, pairwise interactions in three and triple-wise interactions in five animals show no significant difference between stationary and running. In VISrl, firing rates in six out of seven animals show no significant difference between states, while pairwise interactions in four animals and triple-wise interactions in three animals show no significant difference. In VISal, firing rates in five out of seven animals, while pairwise and triple-wise in three animals show no significant interactions. In VISpm, firing rates in all four animals show no significant difference; however, pairwise in two animals show no significant difference. Triple-wise interactions in all animals show significant difference between stationary and running states. In VISam, firing in three out of seven animals, while pairwise two animals show no significant difference between stationary and running states (t-test, $p > 0.05$). Triple-wise interactions in 5 animals doesn't show any significant difference. Data shown here are from brain observatory dataset and the time window is $\Delta = 20$ms.

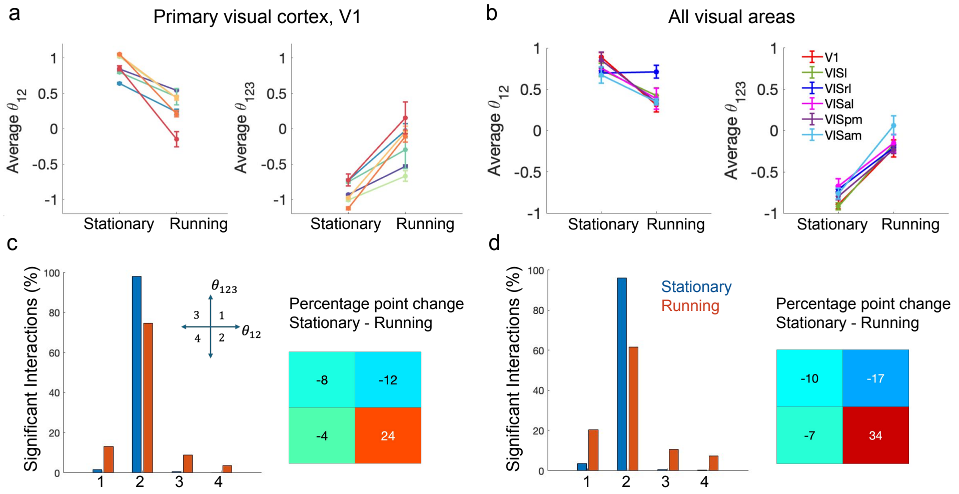


**Supplementary Fig 13.** Pairwise and triple-wise interactions among neurons during stationary and running states in mice. **a** Average pairwise (left) and triple-wise (right) interactions for stationary and running trials of each mouse in primary visual cortex (V1). All animals show significant difference for pairwise and triple-wise interactions between stationary and running states. However, the firing rates in four out of seven animals don't show significant difference (t-test, Suppl. Fig. 15). **b** Pairwise (left) and triple-wise (right) interactions averaged over mice across visual hierarchies. Except for pairwise interactions in VISl and VISrl, all pairwise and triple-wise interactions differed significantly between stationary and running states. However firing rates in VISl, VISrl and VISpm don't show significant difference (t-test, Suppl. Fig 15). Error bars in **a** and **b** represent SEM. The result of each animal in other visual regions are shown in Suppl. Fig. 14 and the firing rates of neurons participating in significant interactions in **a** and **b** are shown in Suppl. Fig. 15. **c** Percentage of significant interactions for stationary and running states in each quadrant of triple-wise versus pairwise plane, for V1 (left) and percentage point change between stationary and running data (right). **d** Percentage of significant interactions for stationary and running states in each quadrant in interaction plane, averaged across all visual areas (left). The percentage point difference between stationary and running data is shown in right panel. The results shown here are from Brain Observatory dataset for bin width 50 ms.

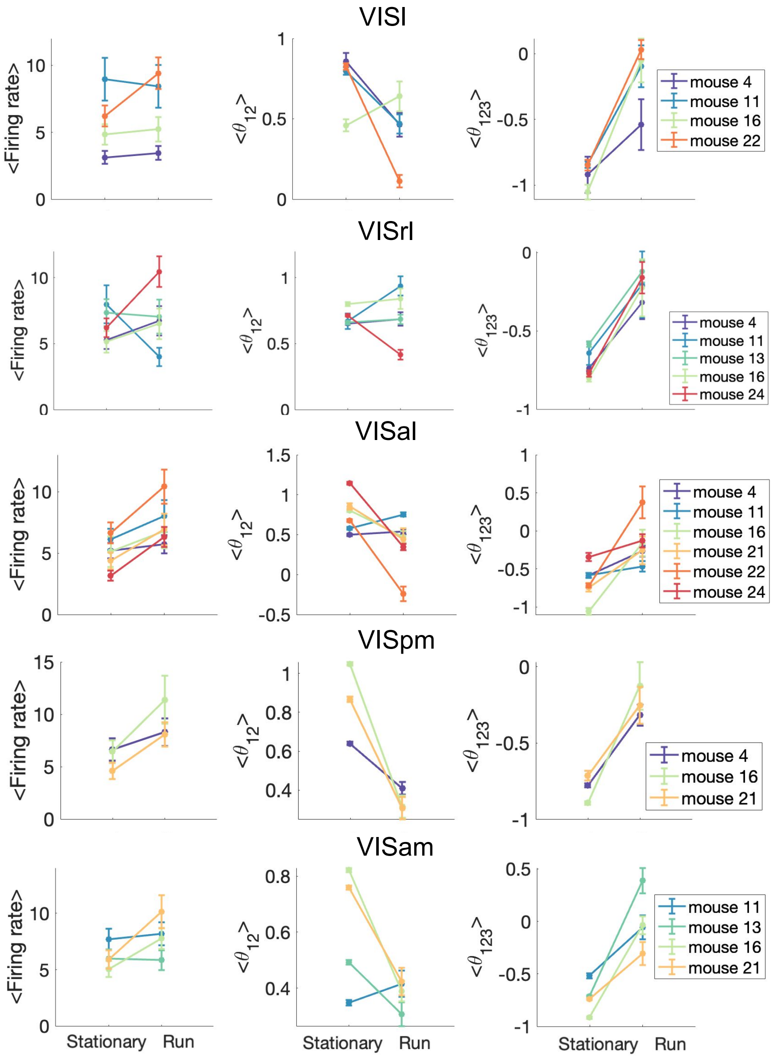


**Supplementary Fig 14.** Firing rates, pairwise and triple-wise interactions between two groups of stationary and running states using bin size 50 ms for Brain Observatory dataset are shown across visual areas. In VISl, firing rates in three out of four mice don't show significant difference between stationary and running states (t-test, $p > 0.05$), while pairwise and triple-wise interactions show significant difference except in one animal. In VISrl, firing rates and pairwise interactions in three out of five mice don't show significant difference between stationary and running states (t-test, $p > 0.05$), but triple-wise interaction show significant difference in all except one animal (t-test, $p = 0.06 > 0.05$). In VISal, firing rates in four out of six mice don't show significant difference between stationary and running groups (t-test, $p > 0.05$). Pairwise and triple-wise interactions were significant across nearly all animals ($p < 0.05$, t-test), with the exception of one animal for pairwise ($p = 0.18$) and two for triple-wise interactions ($p = 0.08$ and 0.055). In VISpm, firing rates in one out of three mice doesn't show significant difference (t-test, $p > 0.05$), while pairwise and triple-wise interactions in all mice show significant difference between states. In VISam, firing rates in three out of five mice don't show significant difference between stationary and running states (t-test, $p > 0.05$).

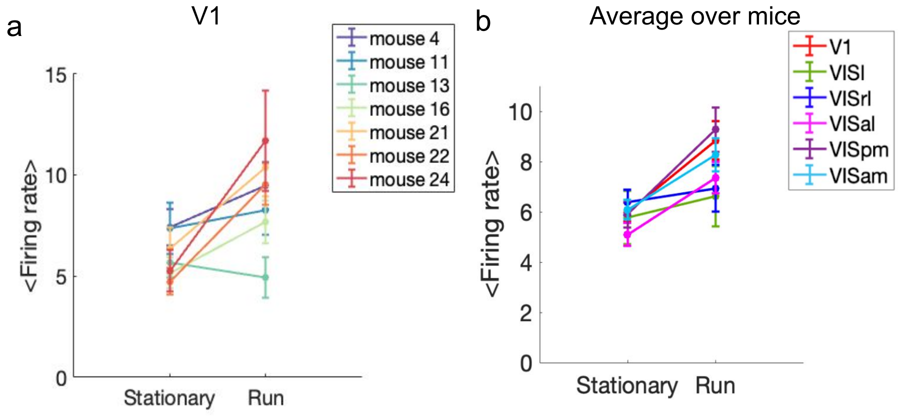


**Supplementary Fig 15.** Firing rates of stationary and running states for each mouse in **a** V1 and **b** across all mice for each visual region. In V1, firing rates in four out of seven mice don't show significant difference between stationary and running states (t-test, $p > 0.05$). **b** Firing rates between two group of stationary and running are not significant in VISl, VISrl and VIspm regions (t-test, $p > 0.05$). The results shown here are from Brain Observatory dataset for bin width 50 ms.

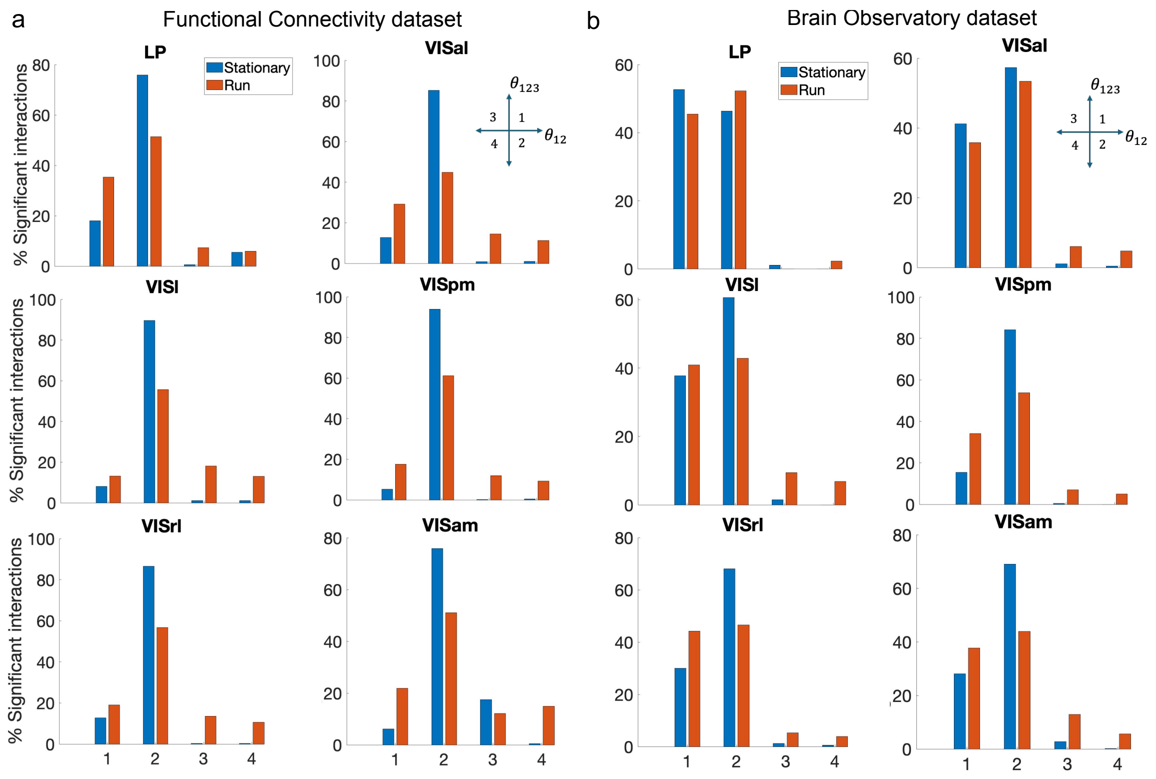


**Supplementary Fig 16.** Percentage of significant interactions in stationary (blue) and running (red) states in each quadrant of pairwise and triple-wise interactions plane for Functional Connectivity dataset (**a**) and Brain Observatory dataset (**b**). The results are shown for visual areas from LP to VISam (LGN doesn't show any significant interactions for stationary and running states). The bin width is 20 ms.

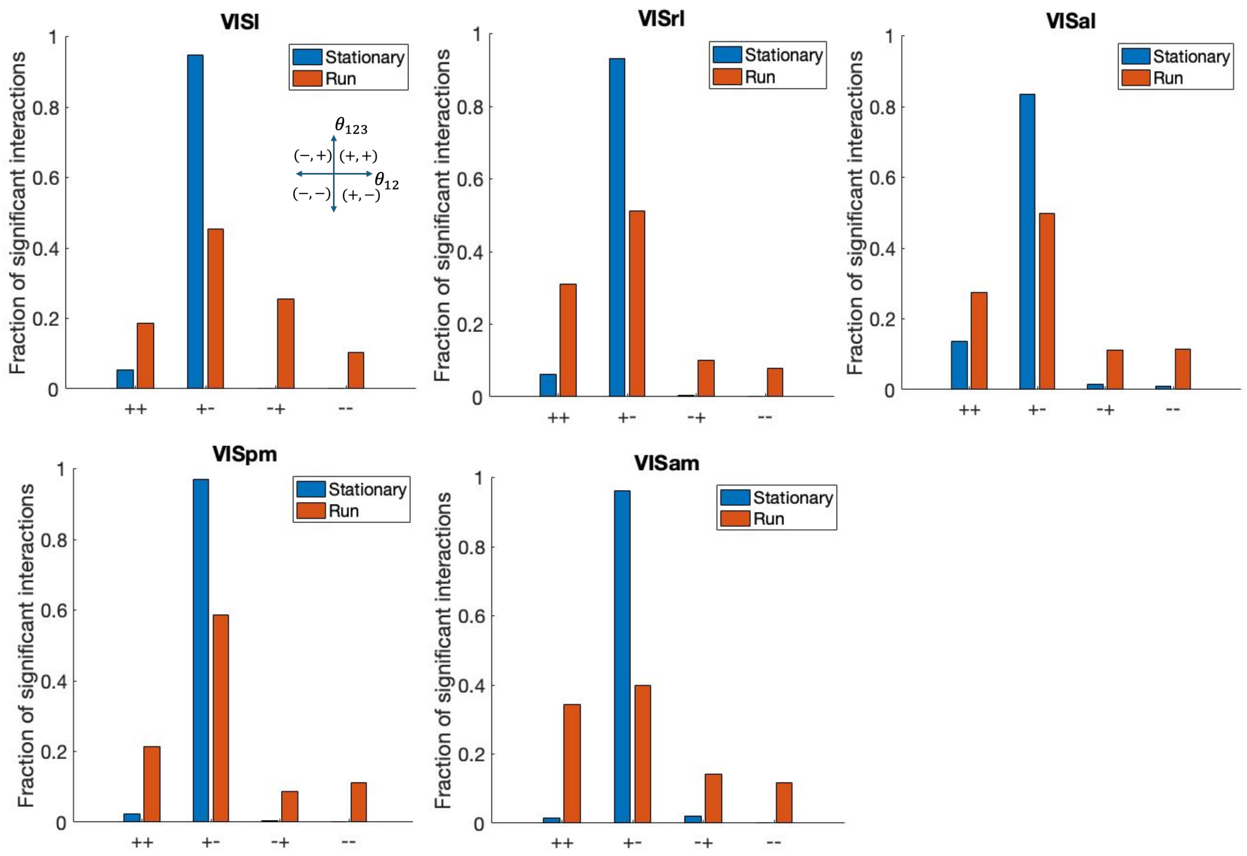


**Supplementary Fig 17.** Fraction of significant interactions in stationary (blue) and running (red) states in each quadrant of pairwise and triple-wise interactions plane for visual regions. The data is from Brain Observatory dataset and the bin width is 50 ms.

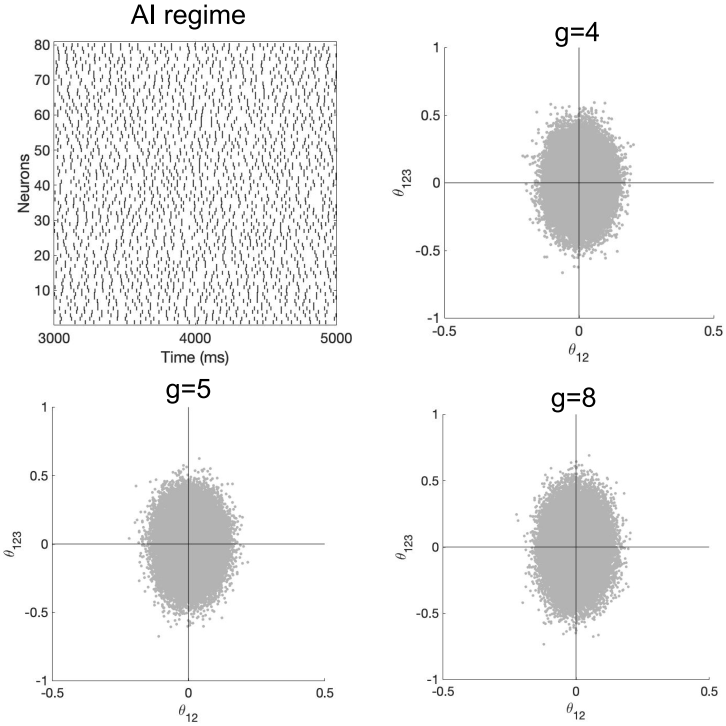


**Supplementary Fig 18.** Balanced random network in asynchronous irregular regime ($J = 0.1$). Top: Left, raster plot of spiking activities is shown for 80 neurons in a random network composed of 800 excitatory and 200 inhibitory neurons. The network shows no significant interactions by varying the balance of excitatory to inhibitory ratios ($g = 4 - 8$). The interactions are shown for a population of 80 neurons and the bin size of 20ms. .

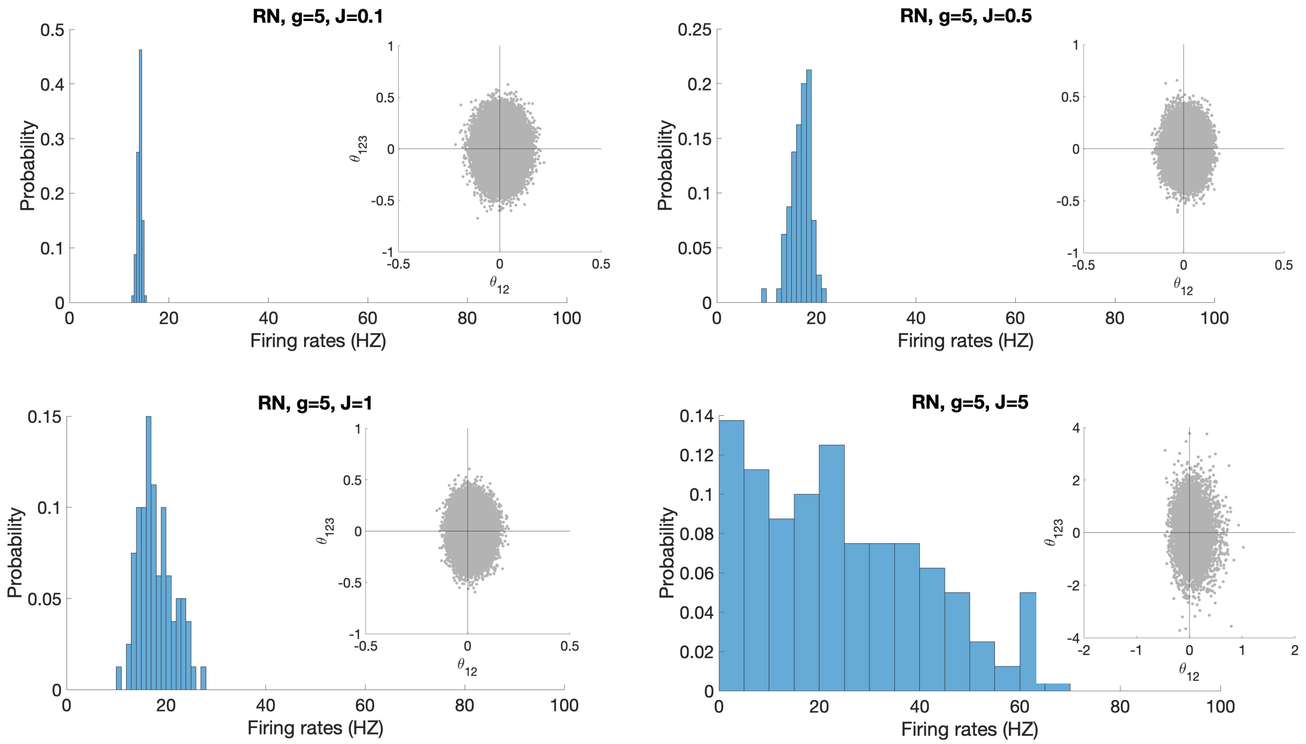


**Supplementary Fig 19.** Balanced random network with different connectivity strength $J$ cannot generate significant interactions. The firing rate histogram and non significant interactions are shown for $J = 0.1-5$ and $g = 5$. As the connectivity strength increases, the firing rate distribution becomes wider.

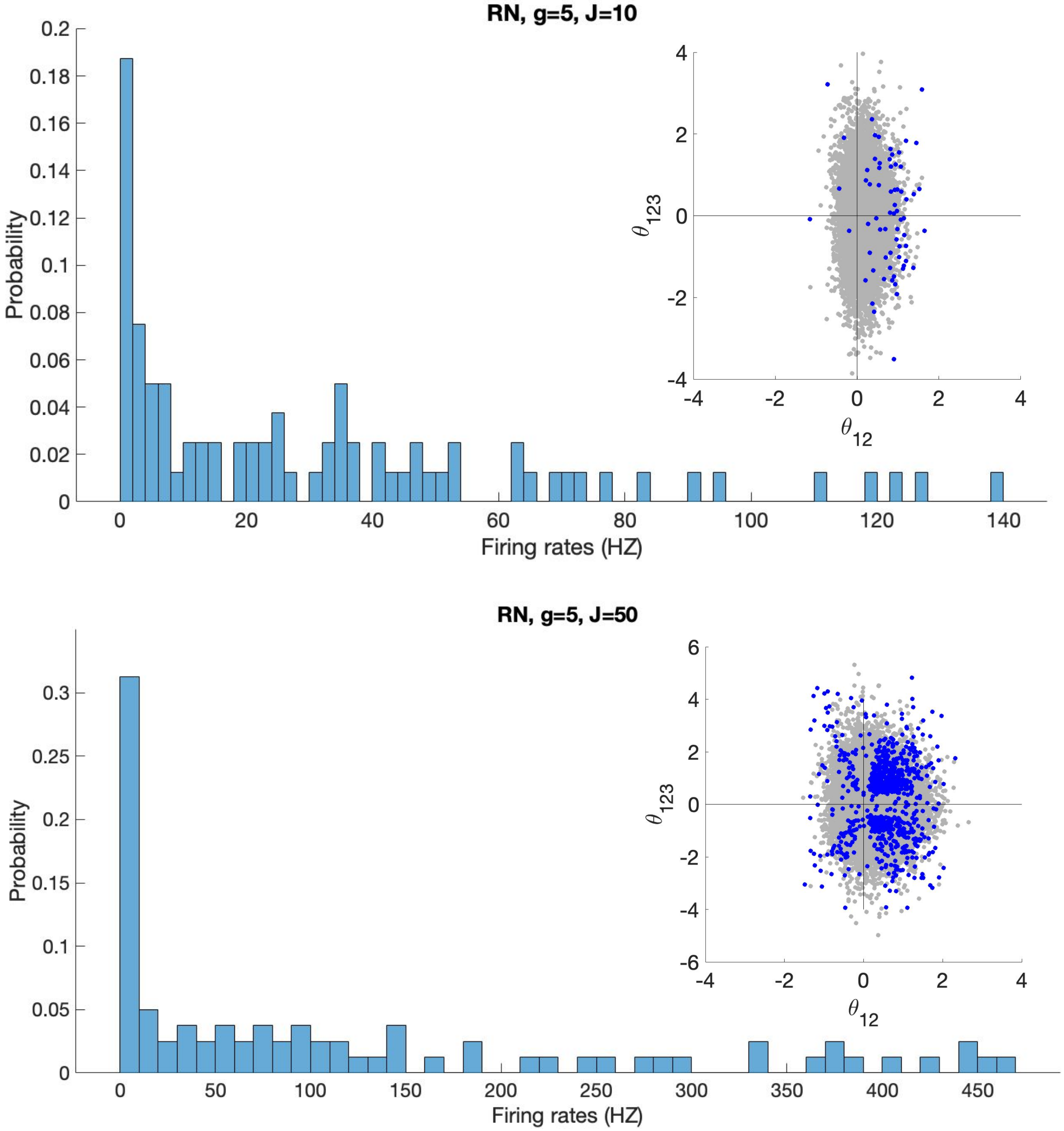


**Supplementary Fig 20.** Balanced random networks for large connectivity strength *J* generate significant interactions, but the range of firing rates are not biologically plausible.

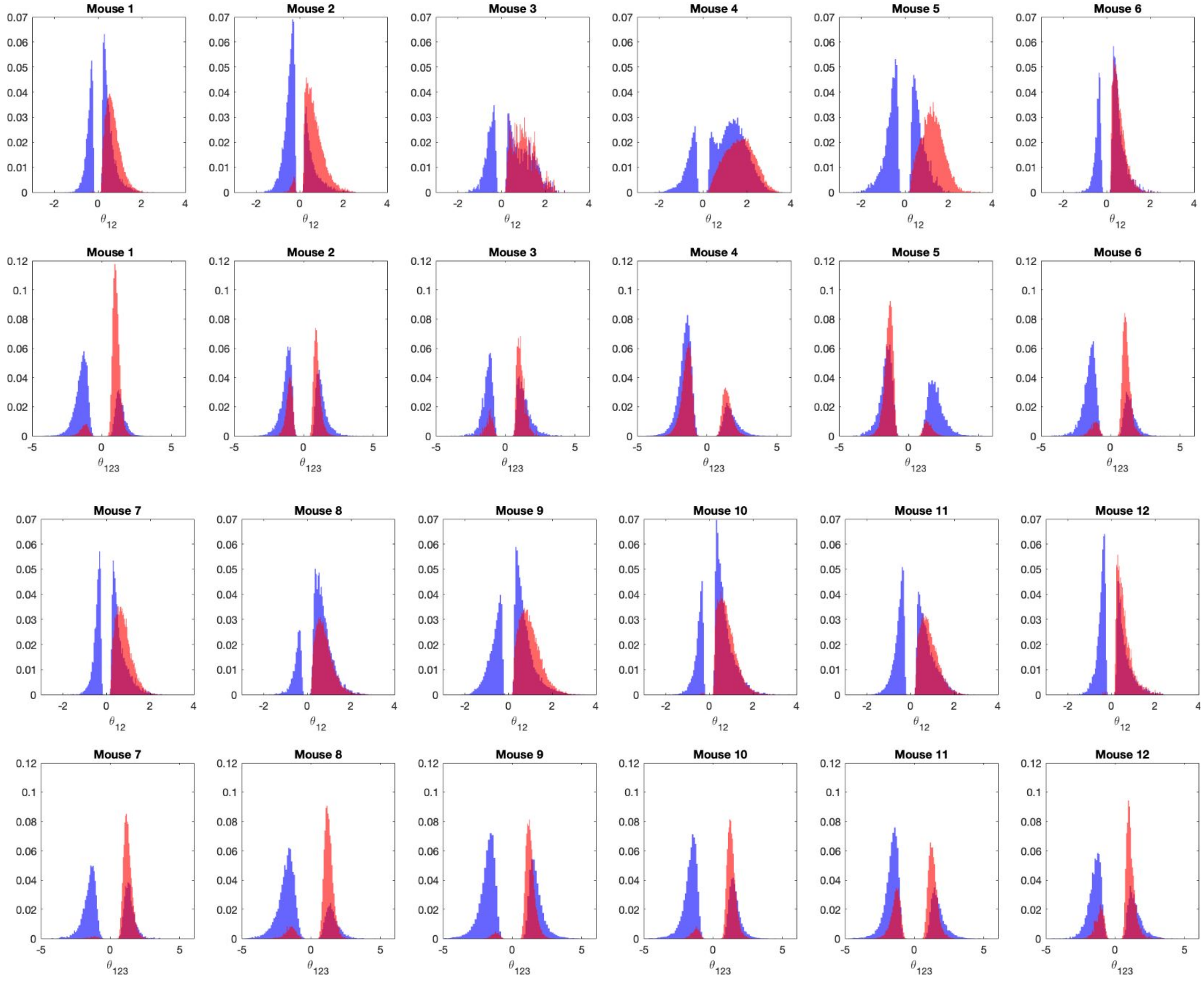


**Supplementary Fig 21.** Distribution of pairwise (top) and triple-wise (bottom) interactions in offsemble (blue) and onsemble (red) groups of neurons in twelve mice. For each mouse, the distribution is plotted over 5 ensembles. The bin size in 80ms.
